# Pleckstrin homology domain-containing serine/threonine kinase plays a crucial role in the survival and phagocytosis of *Entamoeba histolytica*

**DOI:** 10.1101/2025.10.24.684426

**Authors:** Shalini Mishra, Priya Tomar, Alok Bhattacharya, S. Gourinath

## Abstract

Kinases are critical regulators of parasite survival, cytoskeletal dynamics, and endocytic processes. *Entamoeba histolytica* encodes 372 kinases, many of which remain functionally uncharacterized. This study characterizes *EhPHDK*, an AGC-family kinase essential for pseudopod formation, phagocytosis, and pinocytosis. Structural modelling suggests an autoregulatory mechanism via its PH domain, potentially modulating kinase activity. Bio-layer interferometry identified a crucial amino acid sequence facilitating interaction with *Eh*CaBP1, implicating *EhPHDK* in endocytic regulation. Live-cell imaging and immunostaining confirmed its dynamic recruitment to the plasma membrane, colocalizing with *Eh*CaBP1 during phagocytosis initiation and progression. Gene silencing severely impaired parasite viability, reducing growth by 90%, while RBC uptake assays demonstrated its indispensable role in phagocytosis. The regulatory interplay between *EhPHDK* and cytoskeletal components underscores its function in actin-driven processes. Given its divergence from host kinases, *EhPHDK* presents a promising therapeutic target for amebiasis. This study provides novel insights into kinase-mediated cytoskeletal remodelling, advancing our understanding of *E. histolytica* pathogenesis and offering potential avenues for intervention in parasitic survival mechanisms.

**Highlights:**

- *EhPHDK* kinase regulates pseudopod formation, phagocytosis, and pinocytosis.
- Structural modelling suggests PH domain autoregulation of kinase activity.
- *EhPHDK* interacts with EhCaBP1, influencing endocytosis and cytoskeletal dynamics.
- Gene silencing reduces parasite survival by 90%, impairing phagocytosis.
- *EhPHDK* is a potential therapeutic target, diverging from host kinases.

## Introduction

The cause of human amoebic dysentery and a significant portion of morbidity and mortality globally is *E. histolytica.* In terms of the composition of its kinome, *E. histolytica* is unique compared to other protozoan parasites. The parasite has a large signalling network for the sense of its surroundings because it displays more TMKs on average than all protozoan parasites (1,2) These TMKs can transmit signals through downstream effectors that could help parasites adjust rapidly to changes in their host gut environment. Additionally, it has been demonstrated that the amoebic TMKs function as receptors for extracellular signals that can cause basic endocytic processes like the phagocytosis of bacteria and fluid or the invasion of host tissues (3,4). In both situations, an extensive signalling cascade is started, and kinases are crucial to this cascade. However, the molecular mechanism underlying the invasive and endocytic processes’ dependence on Ca^2+^ is still unknown (5,6). The *E. histolytica* genome codes for 36 CaM-like kinases and 27 Ca^2+^ binding proteins (CaBP’s), it is reasonable to assume that these proteins may be working in concert to carry out the Ca^2+^-dependent signalling (7). Another crucial kinase, named C2 domain-containing kinase, belongs to the CaM-like kinases superfamily and is responsible for attracting CaBP1 during trogocytosis and erythrophagocytosis (7,8). Since CaBP1 is in charge of bringing other proteins to the phagocytosis site, its recruitment is crucial for the process to start (9,10). Further, this signalling pathway involves the recruitment of another kinase known as alpha kinase, *Eh*AK1 by CaBP1 in calcium-dependent manner. *Eh*AK1 phosphorylates the G-actin also helps in the recruitment of the Arp2/3 complex at the host cell attachment site (11,12). Interestingly, the actin polymerization at the phagocytosis site cannot begin until the Arp2/3 complex is recruited. It also appears that actin dynamics in vivo depend on *Eh*AK1’s phosphorylation of amoebic G-Actin (13). Lipid modifications, sequential phosphorylation, and other mechanisms can control the sequential recruitment of different kinases in an organized manner, which is necessary for the phagocytosis process to proceed. Moreover, Arp2/3 attracts CaBP3, which aids in actin bundling and phagocytic cup progression. Actin polymers are stabilized by the interaction of *Eh*CaBP5 with the IQ motif of myosin 1B and *Eh*Coactosin with G & F actin (14,15). It was also discovered that only *Entamoeba* species contain the Rho guanine nucleotide exchange factor *Eh*FP10, which has a c-terminal domain that binds and bundles actin filaments (16) After this plasma membrane scission occur and endosome formation take place, then the process of phagocytosis completes.

Recent research has demonstrated that amoeba exhibits distinct ingestion behaviours for both live and dead host cells when living in vivo (17). Due to adherence to host cells, many kinases and phosphatases are activated which facilitate the modification of phosphoinositide’s present at the plasma membrane of *E. histolytica* (18). Numerous biological processes, including membrane trafficking, endocytosis, and cytoskeleton reorganization, are regulated by phosphoinositide’s. PtdIns (4,5)P_2_ are produced in the plasma membrane by type I PIPK, which has recently been demonstrated to be involved in erythrophagocytosis (19). Upon interaction with ligands or host cells (dead or alive), PtdIns (4,5)P_2_ is phosphorylated, resulting in the production of PtdIns (3,4,5)P_3_. The endocytic process’s downstream signalling depends on the synthesis of PtdIns (3,4,5)P_3_ (20). The ingesting of live host cells is known to be facilitated by PtdIns (3,4,5)P_3_, which recruits EhAGCK1, an AGC family kinase specific to trogocytosis. If not, phagocytosing dead host cells involves the recruitment of another kinase from the same family, EhAGCK2 (17). However, in the mammalian system, it is known that PKC regulates actin dynamics by phosphorylating myristoylated alanine-rich C-kinase substrate (MARCKS), and Akt phosphorylates actin in the presence of PtdIns (3,4,5)P_3_ (21). According to the previous findings we can assume that actin polymerization may involve AGCK2, whereas EhAGCK1 may relay signals to form tunnel-like structures during trogocytosis (17).

The AGC kinase superfamily comprises 60 protein kinases, including key members Protein Kinase A (PKA), Protein Kinase G (PKG), and Protein Kinase C (PKC), which are essential regulators of various cellular processes.

The *AmoebaDB* database has identified four Akt-like proteins (*EhAGCK1/EHI_*188930*, EhAGCK2/EHI_*053040, *EhPHDK/EHI_*042150*, and EHI_*199030) within the AGC kinase family in *Entamoeba histolytica*. Among these, *EhAGCK1* and *EhAGCK2* belong to the same subfamily but perform distinct roles in endocytosis. The domain organisation of these two AGC kinase proteins is identical. They have an N-ter plecleckstrin homology domain, a kinase core, and a conserved carboxyl domain in their structure. Their kinase activity is governed by conserved motifs and residues (18). *Eh*AGCK1 (EHI_188930) and *Eh*AGCK2 (EHI_053040) only have two phosphorylation sites, according to the *Insilco* study, but other members of the AGC family require three ordered phosphorylation sites for their optimum function and control. Trogocytosis is the main method used by *E. histolytica* during their invasive phase to destroy living cells. Additionally, investigations state that *Eh*AGCK1 is involved in the fragmentation of living mammalian cells. It does not take part in pinocytosis or other endocytic processes like RBC uptake. *Eh*AGCK2, on the other hand, frequently participates in actin dependent endocytosis. Additionally, it has been suggested that *Eh*AGCK1 and a number of other variables may be crucial for the directional motility of amoebas inside the host intestine. It has been discovered that *Eh*AGCK1 is found close to the plasma membrane inside the trophozoites and is involved in the development of pseudopods, whereas *Eh*AGCK2 is evenly distributed throughout the cytoplasm and also close to the plasma membrane (17). So, far no other kinase’s from AGC superfamily has been identified that plays important role in growth, survivability of the *E. histolytica* parasite. In this paper we have discussed and identified the cruciality of *EhPHDK*/Akt (EHI_042150) protein from AGC superfamily, that interact with CaBP1 plays important role in the growth, survivability and in various endocytic process of parasitic protozoan.

Akt/ PH domain containing protein kinase (PHDK) signalling plays a crucial role in the survival and pathogenesis of various other parasites by modulating apoptotic pathways and cellular responses. In *Leishmania spp.*, activation of Akt prevents apoptosis, promoting intracellular persistence within the host’s macrophages, while its inhibition induces programmed cell death, making it a potential drug target (22). *Trypanosoma cruzi* employs Akt-like kinases to regulate intracellular survival mechanisms, facilitating immune evasion and maintaining its life cycle within host cells (23). Recent studies have explored allosteric inhibitors targeting these kinases as therapeutic interventions. In *Plasmodium spp.*, the role of Akt extends to the mosquito vector, where increased Akt signalling in *Anopheles* mosquitoes reduces malaria parasite prevalence, potentially offering a novel vector-based disease control strategy (24) Additionally, *Schistosoma mansoni* exhibits Akt-mediated cell survivability mechanisms essential for its prolonged infection (25). Given its broad functional significance, Akt kinase remains an intriguing molecular target for parasitic disease intervention, necessitating further exploration of its regulatory pathways and therapeutic implications.

## Materials and Methods

### *E. histolytica* culture and maintenance

Trophozoites of *E. histolytica* strain HM-1: IMSS cl-6 were grown axenically in TYI-S-33 medium at 35.5 °C in 15ml screw-capped round bottom glass tubes (26). To maintain transfectant cell lines as indicated, 10 µg ml^−1^ of either neomycin or hygromycin (Sigma) were added.

### Cloning of various constructs

The CAT gene of the shuttle vector pEhHYG-tet-O-CAT was excised using KpnI and BamHI and the full-length *EhPHDK* gene was inserted in its place in either the sense or the antisense direction. The full-length gene was cloned in BamH1 site in the case of GFP vector resulting in GFP tag on amino terminal of protein. A previous construction of the vector involved cloning the GFP mut3 allele (27) into the distinct BamH1 site of the pExEhNeo plasmid (28).

### Western Blot

For the immunoblot analysis, samples were prepared and separated on 12% SDS-PAGE reducing gels and transferred to nitrocellulose membranes (Amersham) using the semi dry transfer system. Nitrocellulose membranes were then, probed with respective antibodies raised in mice (*EhPHDK* at 1:500) or rabbit (EhCaBP1 at 1:1000; anti GFP at 1:1000) followed by probing with secondary anti-rabbit and anti-mice immunoglobulins conjugated to HRPO (1: 10,000; Sigma). Antigen bands were visualized by ECL reagent enhanced chemiluminescence (Amersham).

### Immunofluorescence staining

*E. histolytica* cells were labelled for immunofluorescent imaging. *E. histolytica* cells were collected by centrifugation and washed before re-suspending in TYI-33 medium. The cells were then transferred onto acetone-cleaned coverslips placed in a petri dish and allowed to adhere for 10 min at 35.5 °C. The culture medium was removed and the cells were fixed with 3.7% pre-warmed paraformaldehyde for 30 min. After fixation, the cells were permeabilized with 0.1% Triton X-100/PBS for 1 min. The fixed cells were then washed with PBS and quenched for 30 min in PBS containing 50 mM NH_4_Cl. The coverslips were blocked with 1% BSA/PBS for 30 min, followed by incubation with primary antibody at 37 °C for 1 h. The coverslips were washed with PBS followed by 1% BSA/PBS before incubation with secondary antibody for 30 min at 37 °C. Antibody dilutions used were: anti-at 1:200, anti-EhCaBP1 at 1:200, anti-rabbit Alexa 488 and 556, anti-mice Alexa556 (Molecular Probes) at 1:300, TRITC-Phalloidin at 1:250. The preparations were further washed with PBS and mounted on a glass slide using DABCO (1,4-diazbicyclo (2,2,2) octane (Sigma) 2.5% in 80% glycerol). The edges of the coverslip were sealed with nail-paint to avoid drying. Confocal images were visualized using an Olympus Fluoview FV1000 laser scanning microscope.

### Time lapse imaging

Amoebic cells expressing *EhPHDK* tagged at its N-terminus with GFP, (NGFP-*EhPHDK*) were plated onto a 35 mm glass bottom dish and allowed to adhere to the dish. RBCs were labelled using DiD, Thermo Fisher Scientific). RBCs were collected by pricking a finger with a needle and collecting the blood in PBS. The cells were washed with PBS twice followed by incubation in PBS containing 10 μM DiD for 10 min at 37°C with intermittent tapping. The reaction was stopped by washing the RBCs with PBS and then the RBCs were kept in ice. Labelled RBC or TRICT-Dextran (Sigma) containing media was added and time-lapse imaging was done using a spinning disk confocal microscope (Nikon A1R, Optics-Plan Apo VC606 oil DIC N2, Camera-Nikon A1, NA-1.4, RI-1.515). The temperature was maintained at 37°C with the help of a chamber provided along with the microscope. The images were captured at 500 ms intervals. The raw images were processed using NIS element 3.20 analysis software.

### Growth Curve of *E. histolytica* trophozoites transfected with Sense and antisense constructs

*E. histolytica* HM1: IMSS trophozoites were transfected with Sense_Toc_ and Anti sense_Toc_vectors. For growth curve studies, the growth of trophozoites in these cell lines with or without induction of tetracycline was measured for 72 hours. The experiment was performed in triplicates in 7 mL tubes. Trophozoites were harvested from a 50 mL culture, counted, and inoculated into each tube. Cell numbers were determined at each time point using hemacytometer.

### Erythrophagocytosis of red blood cells by trophozoites

To quantify the RBC ingested by amoebae, the colorimetric method with some of estimation modifications was used (29). Briefly 107 RBC, washed with PBS and TYI-33, were incubated with 105 amoebae for 20 min at 37°C in 0.2 ml culture medium. The amoebae and erythrocytes were pelleted down; non-engulfed RBC were busted with cold distill water and recentrifuged at 1000 g for 2 min. This step was repeated twice, followed by resuspension in 1 ml formic acid to burst amoeba containing engulfed RBC. Samples were measured against a formic acid blank with a spectrophotometer at 400 nm.

### Interaction study of recombinant purified *EhPHDK* protein and EhCaBP1

A 100 µM *EhPHDK* solution in HEPES was diluted to a final concentration of 20 µM (binding buffer). EhCaBP1 was prepared as a serial dilution (12.5, 25, 50, 100 µM) in HBS-EP running buffer and allowed to bind the *EhPHDK* saturated tips for 600 seconds. Data acquisition was performed using Forte Bio’s Data Acquisition software version 7.1.0.92, and senso grams were analysed using Data Analysis software version 7.1.0.36 on an Octet Red96 instrument (30).

### Statistical analysis

The Statistical comparisons of the data were performed using a two-way ANOVA test by GraphPad Prism Software (version 8.0.2). P values of <0.05 were considered statistically significant. Pearson’s Correlation Coefficient were obtained either using Olympus Fluoview FV1000 software.

## RESULTS

### Evolutionary conservation and structural modelling reveal domain architecture of *Eh*PHDK macromolecule

To evaluate *Eh*PHDK as a biologically active macromolecule, we performed a comprehensive sequence and structural analysis to uncover its conserved domain architecture and evolutionary features. Sequence study reveal that *EhPHDK* shows sequence similarity with other Akt isoforms from different organisms including various parasites, yeast, fungi, Bos taros, Staphylococcus aureus and human. *EhPHDK* protein shows maximum 45 % sequence identity with Dictyostelium discoideum serine/threonine kinase. Identity ranged from 33% with (Leishmania infantum, Leishmania panamensis, Leishmania donovani) to 42% with serine/threonine protein kinase from human (S Fig1A and S Fig1B.). The multiple sequence alignment and 3D structure predicted through Alpha Fold software shows that all the PH domain containing protein kinase possess three functional domains i.e., PH domain, kinase core and a C terminal domain (Fig.1A). The PH domain interact with very high affinity with phosphatidyl inositol 3,4,5 triphosphate that shows it’s a membrane interacting protein (17). The alignment result also proves that only the kinase core is highly conserved among all the organisms and especially among the group of parasites. While the sequences from other two domains show high range of variability, that makes these Akt isoforms distinct from each other. The Alpha fold model suggests that the PH domain contains eight antiparallel beta sheet and one long alpha helix (Fig.1. B. i) that attach with a kinase core via long stretch of 18 amino acid residues. The kinase core consists of seven anti-parallel beta sheet, eight alpha helices and highly disordered loops. The active site or polypeptide substrate binding site also known as activation loop/A loop lie within a kinase core (residue 237-249), (Fig.1. B. ii & S. Fig.2). The long C terminal region (Fig 1 B.i) seems to be very flexible and wrap around the kinase domain. Structural modelling further indicates that this C-terminal segment, together with the PH domain, sterically occludes the kinase core’s active site, suggesting an autoregulatory mechanism whereby conformational changes modulate substrate access. Notably, the pronounced structural divergence of the PH and C-terminal domains from those of other Akt homologs highlights evolutionary tuning for parasite-specific regulation.

**Figure 1.**
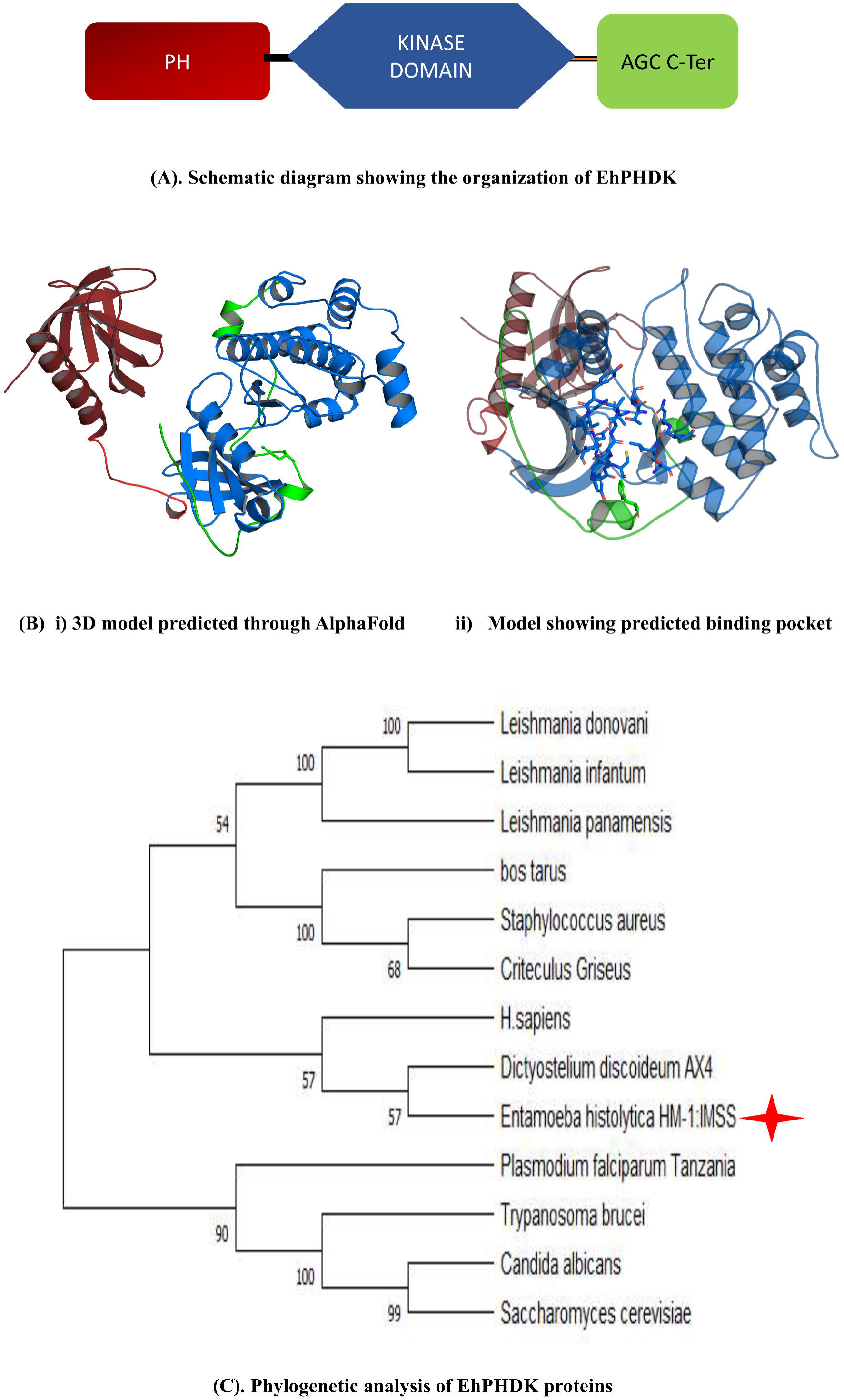
Domain architecture of *EhPHDK* and their phylogenetic analysis. **(A)**. Schematic diagram showing the organization of *EhPHDK* (PH domain (red), kinase core (blue) and C-terminal domain(green). (B) 3-D structure of full length *EhPHDK* using Alpha-Fold software (left) and model showing ATP/peptide binding pocket (right). **(C)** Phylogenetic analysis of *EhPHDK* proteins. Phylogenetic tree was constructed using an iterative neighbor-joining algorithm of MEGA 7.0 software. Bootstrap values (as percentages) are shown at each node.

**Figure 2.**
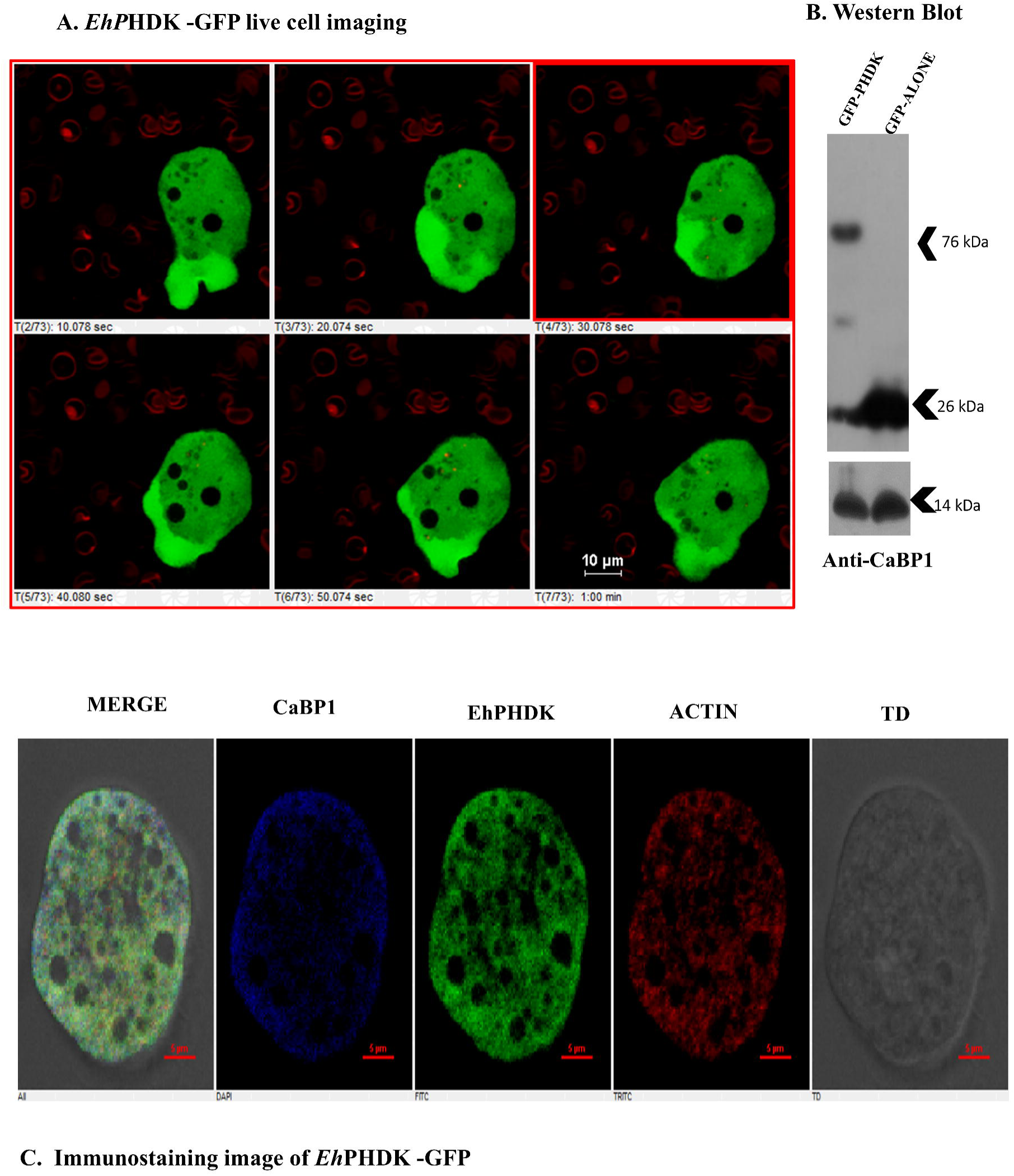
GFP-*EhPHDK* dynamics in moving trophozoites are displayed in a live-cell imaging montage. (A) The montage shows a time series of chosen fluorescent frames of motile trophozoites that are expressing GFP-*EhPHDK*. Pseudopods with higher GFP-*EhPHDK* fluorescence intensity (in presence of RBC). (B) Western blot detection of the GFP-*EhPHDK* in the parasite lysate. 150 µg of the lysate from *E. histolytica* cells expressing GFP-*EhPHDK* and GFP alone, cultured in the presence of 30 µg/ml G418 was loaded in each lane and probed with anti-GFP antibody. In GFP-*EhPHDK* expressing cells, an anti-GFP antibody identified a 50 kDa protein (*EhPHDK* + GFP) band when compared to a 26 kDa band of GFP alone in control cells. Anti-EhCaBP1 antibody was used as an equivalent loading control. (C) Immunolocalization of *EhPHDK* in fixed trophozoites (without RBC) demonstrating its cellular localization and beneath the membrane at specific location.

### Cellular localization and dynamics of *EhPHDK* in proliferating trophozoites

Phagocytosis is a key mechanism responsible for tissue invasion and pathogenesis of this parasite which causes amoebic dysentery. To look into the cellular dynamics, localization, and their role in pseudopod formation in proliferating trophozoites, N-Ter GFP tagged *EhPHDK* genes was firstly cloned in pEh-neo-GFP vector and transfected in wild-type HM1: IMSS strain of *Entamoeba histolytica.* The expression of N Ter-GFP tagged *EhPHDK* protein (50kDa *EhPHDK*+26 kDa GFP) was confirmed through western blot analysis by using an anti-GFP antibody (Fig2 B). We carried out live cell imaging of GFP-fused *EhPHDK* transformant cells, which revealed *EhPHDK* found ubiquitously in the cytoplasm in proliferating trophozoites. Also, the expression was found to be enriched along with the pseudopod formation and during the membrane invagination phenomenon (Fig2A). To further investigate the cellular dispersal of *EhPHDK* in normal homeostasis, an immunostaining study was conducted in the absence of RBCs. This study confirmed that *EhPHDK* is primarily a cytosolic protein (Fig2C). Additionally, it was found to become enriched beneath the membrane at specific sites (in the presence of RBCs), leading to overexpression of this protein in cup-like projections and membrane protrusions Additionally, we conducted fluorescence intensity profiling using live-cell imaging and immunostaining, comparing it against controls (WT trophozoites and neo-GFP transformants). This experiment aimed to determine whether cytosolic GFP accumulates or is excluded from pseudopods (S. Fig.3 A, B, C). By measuring fluorescence intensity from the budding pseudopod (S/ Start) to the opposite side (E/ End), we found GFP-tagged *EhPHDK* was enriched beneath the pseudopod, confirming its preferential localization within these dynamic structures. In contrast, neo-GFP showed uniform cytoplasmic distribution, and wild-type HM1 cells displayed negligible fluorescence, confirming no intrinsic GFP expression.

**Figure 3.**
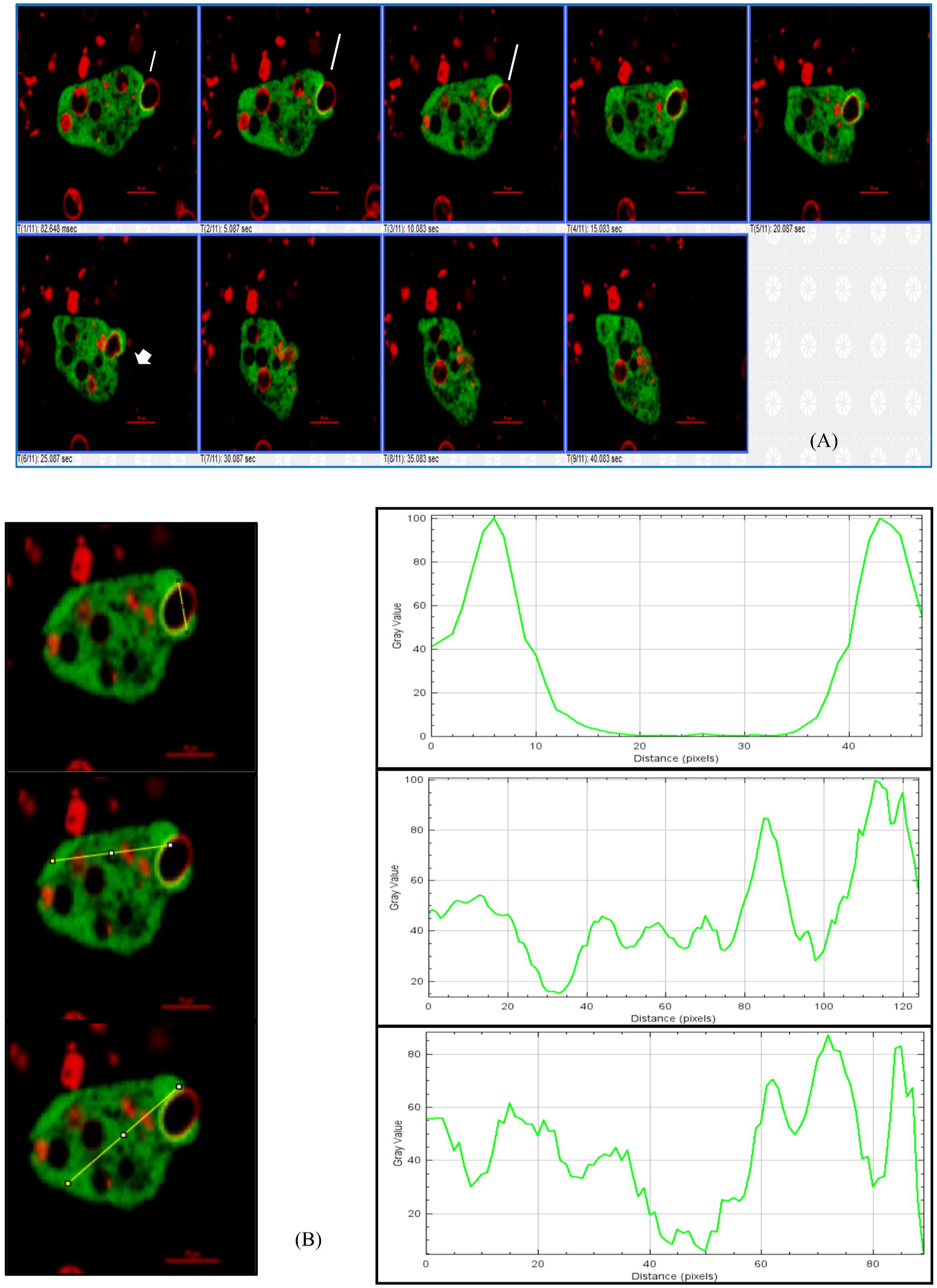
Time-lapse imaging of GFP-*EhPHDK* during erythrophagocytosis. (A) The montage displays time-lapse pictures of the phagocytosis of RBCs by an amoeba cell expressing GFP-*EhPHDK*. The DiD stain was applied to RBCs. The represented figure demonstrates *EhPHDK* active involvement in the phagocytic cup’s creation and closure, which is highlighted by white arrowheads. At the RBC attachment site, GFP-*EhPHDK* quickly gathered and left as the phagosome closed. (10 µm scale bar). (B) Graph displaying the fluorescence intensity of several cell types expressing *EhPHDK*-GFP. Using ImageJ software, fluorescent signal intensity was assessed in three randomly chosen locations inside the phagocytic cup, cytosol, and plasma membrane.

This direct comparison ensures that *EhPHDK* enrichment is not an artifact of GFP expression but reflects its functional role in pseudopod dynamics. By integrating both fluorescence intensity profiling and immunostaining, this study provides strong evidence that *EhPHDK* is functionally involved in pseudopod formation, distinct from general GFP expression patterns. These findings enhance our understanding of the molecular dynamics of cytoskeletal remodelling in amoebic trophozoites and may contribute to further studies on *E. histolytica* motility and invasion.

The image panel from the live cell imaging and immunostaining proves that *EhPHDK* is initially present in the cytoplasm but during the host cell attachment and phagocytosis phenomenon *EhPHDK* protein actively colocalise with actin and helps in the formation of pseudopod (S. Fig.4A). The abundance of the protein in pseudopods in comparison with the cytoplasm of the cells was further validated by intensity profiling by choosing a micrograph from the random pseudopod-like structure (S. Fig.4B). Similar *EhPHDK* protein localization and expression pattern were also observed when we carried out immunostaining of wild-type cells with an *EhPHDK* antibody (1:1500) (S. Fig.5 A & B). The results predicts that GFP fused protein shows a similar localization pattern as the native protein shows i.e., it mainly presents in cytoplasm but during the attachment, cup formation and membrane protrusion it actively colocalise with actin and shows the enrichment underneath the membrane at pseudopods.

**Figure 4.**
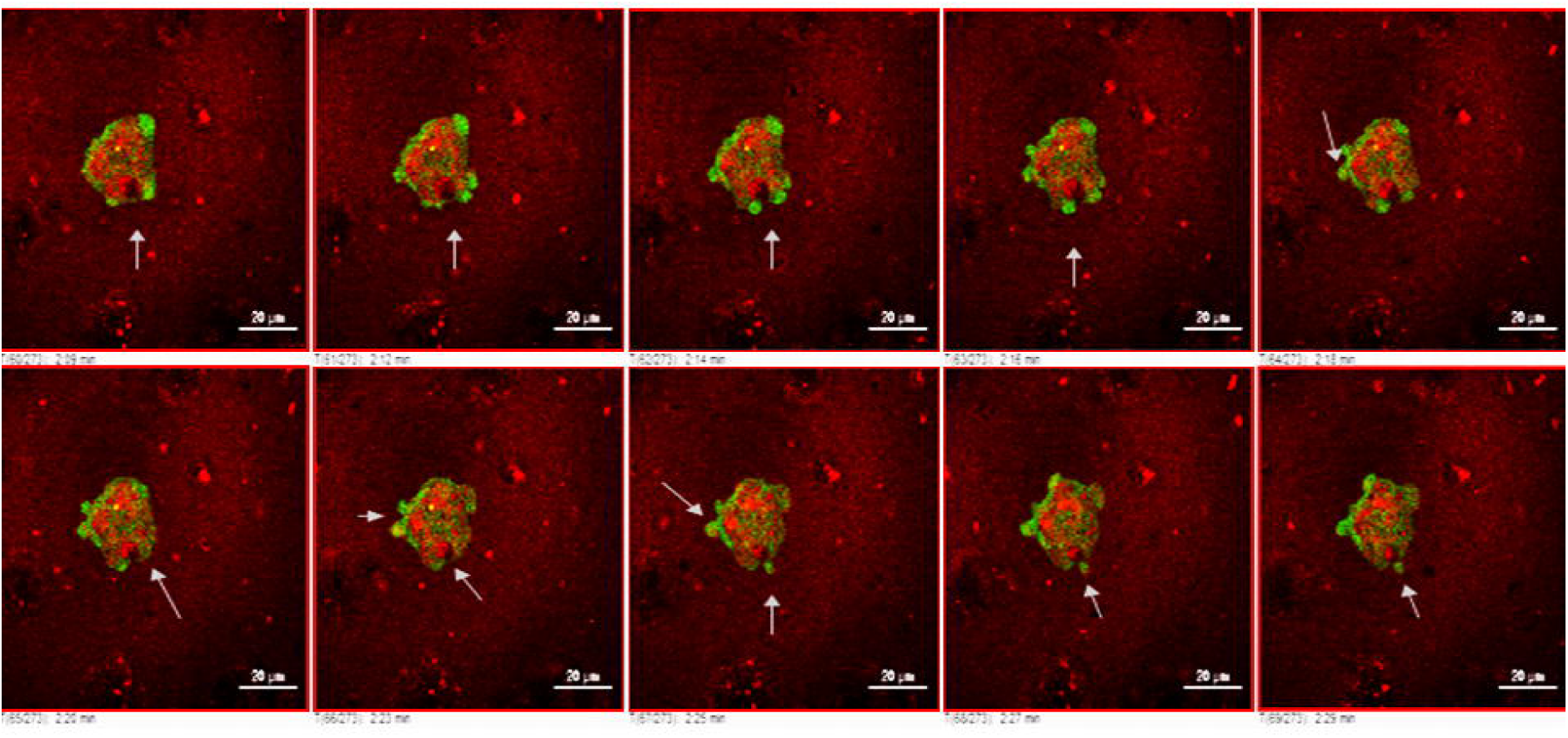
Active localization of EhPHDK during fluid-phase micropinocytosis: GFP-*EhPHDK* time-lapse imaging during fluid-phase micropinocytosis. The media was provided with TRITC Dextran, which gave it a red fluorescence. Red fluorescence within internalized vesicles enriched with GFP served as evidence for the participation of GFP *EhPHDK* in fluid-phase micropinocytosis (marked with arrow).

**Figure 5.**
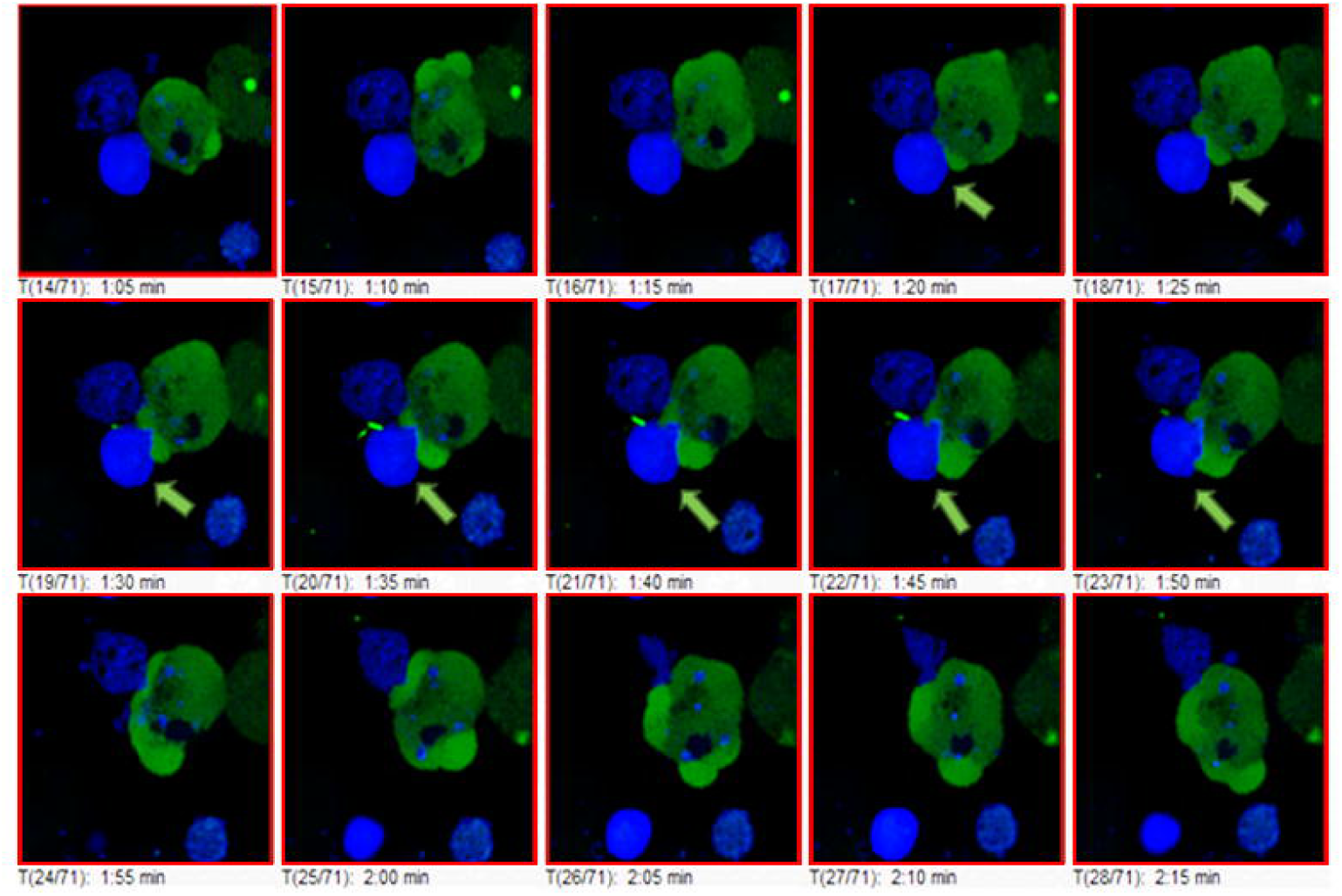
Absence of EhPHDK Recruitment during Trogocytosis: The montage showing the *Invivo* localization of GFP-*EhPHDK* during the uptake of CHO cells, where CHO cells are stained with (CMAC) Cell tracker blue dye. No trogocytosis event was observed in any cells. While the complete phagocytosis of CHO cells occurs.

### *EhPHDK* actively participate in pseudopod formation and phagocytosis

Tissue invasion phenomenon involves three steps i.e., adherence, cytolysis, and phagocytosis of dead cells or trogocytosis of live cells. Adherence leads to the activation of several kinases and phosphatases that help in the modification of phosphoinositide present on the plasma membrane of an amoeba (31). The elevated calcium ion and phosphoinositide concentration in the cell helps in the recruitment of various proteins at the attachment site and regulates the cytoskeleton reorganization that results in pseudopod formation (32)

The enrichment of *EhPHDK* was observed at pseudopods in live cell imaging of proliferating trophozoites expressing GFP-*EhPHDK*, indicating *EhPHDK* actively participates in pseudopods. At the moving pseudopods edge of the amoeba, the enrichment of protein was detected that are highlighted by the arrow ahead in the figures (Fig3 A). The further enrichment analysis was quantified by line intensity plot by selecting three random regions inside the cell. The graph shows the relative fluorescence intensity of GFP-*EhPHDK* in cup projection vs cytosol and plasma membrane along the drawn line using Image J software. (Fig 3B). Although the trogocytosis phenomenon in RBCs has been documented, we were not able to identify it in these tests using microscopy (33,34). This may be possible because trogocytosis is favoured for the flat, large, live host cells, whereas phagocytosis may be preferred for the small, disc-shaped RBCs (17).

### *EhPHDK* involved in fluid-phase micropinocytosis but not in trogocytosis

We further explore the role of *EhPHDK* in other endocytic process like micropinocytosis and trogocytosis. The majority of the nutrients that amoebic cells in culture and the host intestine absorb come via pinocytosis or micropinocytosis. Time-dependent TRITC dextran uptake was used to investigate the pinocytosis event of *E. histolytica* trophozoites. The medium was supplemented with TRITC Dextran, which produced a red fluorescence. The appearance of red fluorescence within internalised vesicles enriched with GFP established the participation of *EhPHDK* in fluid phase micropinocytosis. Also, GFP-*EhPHDK* quickly dissociated after the pinocytic vesicle was closed. A single micropinocytosis event typically takes 6-7 seconds to complete. (Fig 4.)/ (S. vedio2).

Amoebic cells ingest live mammalian cells using trogocytosis (35). This type of endocytosis becomes crucial throughout the parasite’s invasive phase. The catastrophic destruction of CHO cells by trophozoites resembles the devastation of host cells after invasion by the parasite (36). Therefore, with live cell imaging, the trogocytosis of live CHO cells was also examined by proliferating trophozoites. A time-lapse montage of GFP-*EhPHDK*-expressing cells revealed that GFP-*EhPHDK* is attracted to the location where the trophozoite establishes contact with the CHO cell before proceeding through the tunnel and engulfing the entire cell. So, unlike *Eh*AGCK1 and *Eh*AGCK2, which have already documented to be involved in the trogocytosis process (16), *EhPHDK* doesn’t show any participation in the nibbling of live cells.

Using bigger sized mammalian Chinese hamster ovary (CHO) cells also allowed us to more clearly see where *EhPHDK* was located at the phagocytic cup. Trophozoite were observed during the different stages of phagocytosis when they were treated with CHO cells that had been stained with a cell tracker dye (Fig. 5/ S. video 3). The analysis of phagocytic cups and phagosomes supports the idea that *EhPHDK* remains at the site of phagocytosis until membrane closure.

### Active colocalization of *EhPHDK* with *Eh*CaBP1 during the initiation and progression step of phagocytosis

The bulk of endocytic processes, including trogocytosis, phagocytosis, and micropinocytosis, require phosphoinositide’s and the actin cytoskeleton. The signalling processes that link these two components together in *E. histolytica* are not fully understood. The other two member of AGC superfamily i.e., AGCK1 and AGCK2 already known to interact with very high affinity with phosphoinositide’s and regulate the cytoskeleton reorganisation. According to previous findings, it has been shown that *Eh*CaBP1, *Eh*Coactosin, *Eh*AK, and *Eh*C2PK play a part in the onset of phagocytosis and the formation of phagocytic cups in *E. histolytica* (5,15,6,7). However, only EhCaBP3 was found in mature phagosomes, and neither EhCaBP1 nor EhC2PK were found in cups that had just closed before scission from the membrane.

To better understand the mechanistic details of *EhPHDK* with *Eh*CaBP1 that has been identified as being a part of the initiation and progression step of phagocytosis pathway in amoeba, immunostaining was done and the degree of co-localization during phagocytosis was also measured using (PCC) Pearson correlation coefficient. Overexpressed N-ter GFP tagged *EhPHDK* localisation inside the cells indicates that it participates actively in the phagocytic cup initiation, progression, and closure as well as being present at the cup tip when the process is complete. In all three stages, *EhPHDK* and *Eh*CaBP1 actively co-localize during the onset and development of the pseudopod formation phenomena (Fig.6A). The variation in GFP_PHDK intensity along the arrow is depicted in the micrograph (Fig.6B & C). The trophozoite-formed pseudopod’s intensity profile demonstrates enrichment along the line of analysis and also demonstrates its localization with *Eh*CaBP1 (Pearson’s correlation: 0.7849) (Fig.6 D). Also, in terms of its interaction with phagocytic machinery, it thus appears that *EhPHDK* operates similarly to some other known phagocytic markers such as *Eh*ARPC1 and *Eh*AK1. Both of these molecules escape from phagosomes either in advance of or right after scission (10 & 11). The findings also suggests that *EhPHDK* may recruited at the site of phagocytosis by *Eh*CaBP1, and this interaction is transient i.e., *Eh*CaBP1 remains with this kinase during nucleation and progression step and leaves during the cup closure phenomenon.

**Figure 6.**
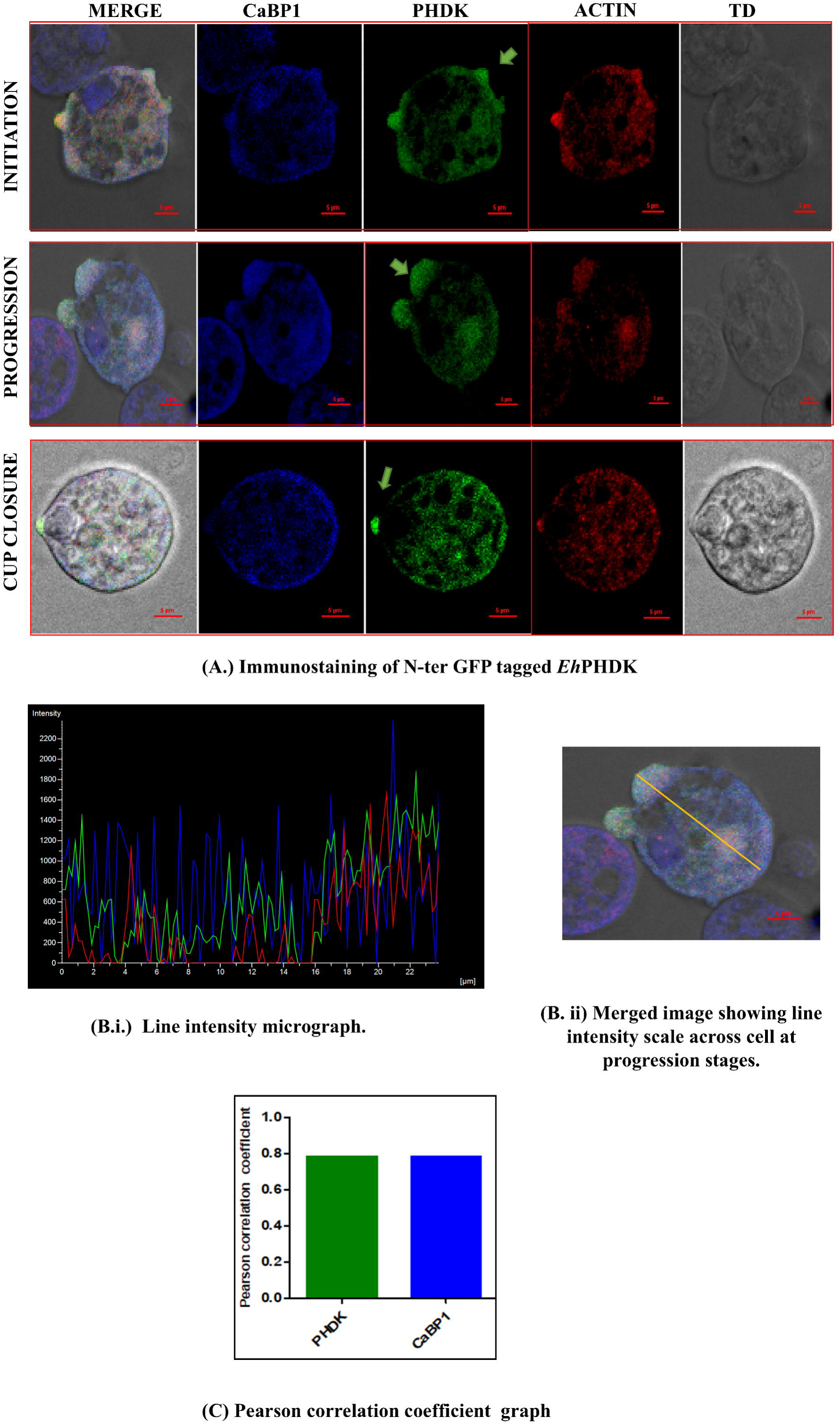
Colocalization of *EhPHDK*/Akt with EhCaBP1. **(A.)** Immunostaining of N-ter GFP tagged *EhPHDK* suggest that it actively involved during the phagocytic cup initiation, progression as well as it is also present at the tip during the cup closure and leave the site by the process complete. Among all the three stages, *EhPHDK* actively co-localise with EhCaBP1 during initiation and progression phenomenon of pseudopod formation. **(B.)** The intensity of GFP_PHDK along the arrow shown in micrograph at different points. The intensity profile at pseudopod formed by the trophozoite show enrichment along the line of analysis and also show its localization with EhCaBP1 (Pearson’s correlation 0.784905). **(C)** Pearson correlation coefficient value was calculated using NIS element software by selecting area enclosing the phagocytosis cup, at the base and also the tip of endocytic cups from different trophozoites. Statistical analysis was performed by using graph pad prism software.

### Macromolecular interface-identification of *EhPHDK* as a binding partner of *Eh*CaBP1

Using a fixed cell immunofluorescence labelling technique and an anti-GFP antibody, we were able to confirm that *EhPHDK* was involved in the various stages of phagocytosis and showed active co-localization with EhCaBP1 and actin. To further understand if these two proteins are directly interacting, Bio-layer interferometry (BLI) experiment was performed between *EhPHDK* and *Eh*CaBP1. The initial interaction investigation was monitored between purified recombinant *EhPHDK* and *Eh*CaBP1 proteins. For, which *EhPHDK* was immobilized onto a Ni-NTA chip and serially diluted *Eh*CaBP1 was employed as an analyte. Both the proteins directly interact with each other. *Eh*CaBP1 shows micromolar binding affinity (i.e. 32µM) with *EhPHDK*/Akt kinase (Fig 7 A). Further, to determine the exact region of *EhPHDK* protein that act as binding sites for *Eh*CaBP1, we have screened and selected a few peptide sequences of *EhPHDK* using imed software (http://imed.med.ucm.es/Tools/antigenic.pl). The three-dimensional domain organization and multiple sequence alignment results stated above show that *EhPHDK* possesses three functional domains and has conserved regulatory sequences that may play an important role in regulating protein activity. The PH domain is connected to a kinase core by a hinge region that possesses IQ motif like amino acid residues (**I**YKSKITTET**V**T**Q**K**D**) which may serve as a binding site for various calcium-binding proteins. The length of the IQ motif is roughly 25 amino acids, and it is found throughout nature. The motif has a consensus sequence [**I**,L,**V**]**Q**xxxRGxxx[R,**K**], which can bind calmodulin without utilizing Ca^2+^ (37).

**Figure 7.**
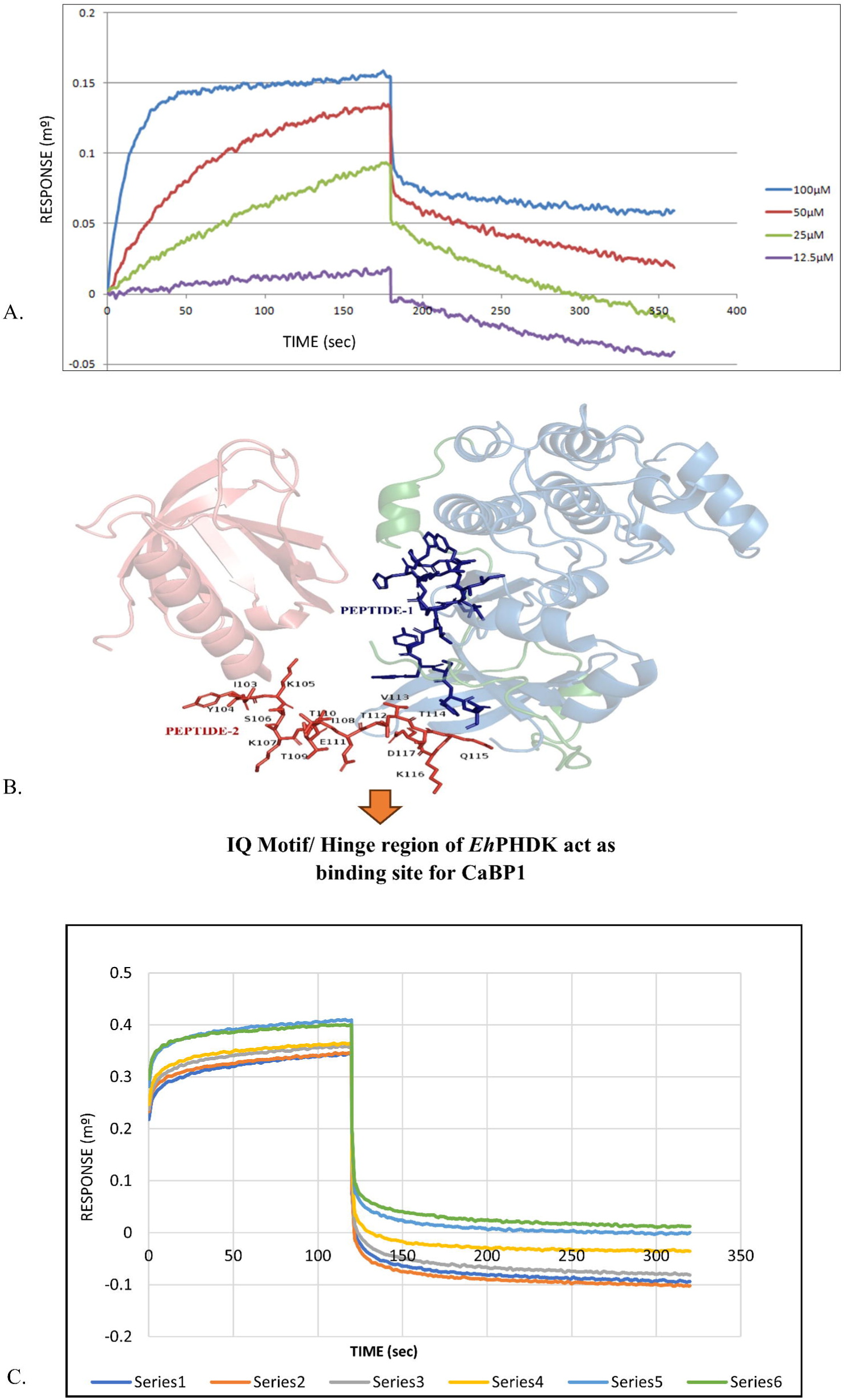
Direct Binding of EhPHDK to EhCaBP1 via IQ-Like motif identified by Bio-layer interferometry. (**A**). BLI study reveal that *EhPHDK* shows interaction with *Eh*CaBP1. The response showing the interaction between *EhPHDK* full-length protein (overlayed onto a Ni-NTA chip) and increasing concentration of recombinant *Eh*CaBP1 (12.5, 25, 50, 100 µM) shows K_d_ of 32 µM. **(B).** Three-dimensional structure predicted through AlphaFold software shows the hinge region (IYKSKITTETVTQKD) lies between the PH domain and a kinase core (highlighted with red colour) serve as binding site for *Eh*CaBP1. **(C).** Bio-layer interferometry graph showing the binding affinity of *EhPHDK* Peptide2 for *Eh*CaBP1(overlayed on Ni-NTA chip. The colours represent different peptide 2 concentrations, ranging from (300 µM-550 µM /series1-series6) exhibits high affinity for *Eh*CaBP1. The Kd value is 243 µM calculated from graph.

Thus, two peptide sequences named Peptide 1 (ILSKLHHPFLVNLYYSFQ) present at the surface of the kinase core and Peptide 2 (**I**YKSKITTET**V**T**Q**K**D**), a sequence present at the hinge region (lies between the PH domain and kinase core) were selected for the protein-peptide binding studies. The *Eh*CaBP1 was immobilized on Ni-NTA chip and both peptides were employed in different concentrations ranging from 50µM to 550 µM. The BLI graph is displayed in Fig. 7. C shows the hues representing different Peptide 2 concentrations (i.e., 300, 350, 400,450, 500, 550 µM/ series 1-6), shows high binding affinity of *EhPHDK* Peptide 2 for *Eh*CaBP1 i.e. 243 µM.

The findings revealed that peptide 2 (hinge region) had the highest binding response with *Eh*CaBP1 in comparison to peptide 1. Peptide 1 doesn’t show any kind of interaction (S. Fig.6), which also supports the fact that Peptide 2 (hinge region) may mimic IQ motif-like sequences that may serve as binding sites for various calcium-binding proteins to regulate the function of kinase and phagocytosis phenomenon.

### *EhPHDK* colocalised with actin and Coactosin

Thus far, we have demonstrated that the *EhPHDK* protein colocalizes with both the phagocytic and pinocytic cups in immunofluorescence experiments using GFP-*EhPHDK* overexpressed cells (Figure 3 & 4). Co-localization of *EhPHDK* with *Eh*CaBP1 and actin was observed from cup initiation to vesicle scission (Fig 6 A). Actin is more locally localized in the base of the cup, whereas *EhPHDK* and *Eh*CaBP1 colocalize at the proximal end of progressing pseudopods. All three of them colocalized incredibly well as the cup was about to close. Like *Eh*CaBP1, Coactosin protein is also a known phagocytic marker that contributes to the initiation and development of phagocytic cups in *E. histolytica* (28).

In our immunostaining study *EhPHDK* was observed to colocalize with *Eh*Coactosin at initiation stage of phagocytosis (S. Fig7). Its localization with *Eh*Coactosin (Pearson’s correlation 0.7075) and enrichment along the line of analysis are both evident in the intensity profile at the budding pseudopod formed by the trophozoite.

### Down regulation of *EhPHDK* gene significantly declines the survivability and phagocytosis process of *Entamoeba histolytica*

In order to investigate the biological importance of *EhPHDK* in several cellular processes of *E. histolytica*, we have transfected *E. histolytica* HM1: IMSS strain trophozoites with tetracycline-inducible vectors Tet-O-CAT that have PHDK gene cloned in both the sense (S_Toc) and antisense (AS_Toc) directions (Fig 8 A). By performing reverse transcription PCR on the corresponding cDNA, (Fig. 8 B) and with western blot analysis the precise gene suppression was verified (S. Fig 8). According to the relative mRNA expression results, the *EhPHDK* gene has been specifically silenced. And by over-expressing the gene in an antisense orientation in a tetracycline-inducible manner, it was possible to reduce the expression of *EhPHDK* in proliferating trophozoites.

**Figure 8.**
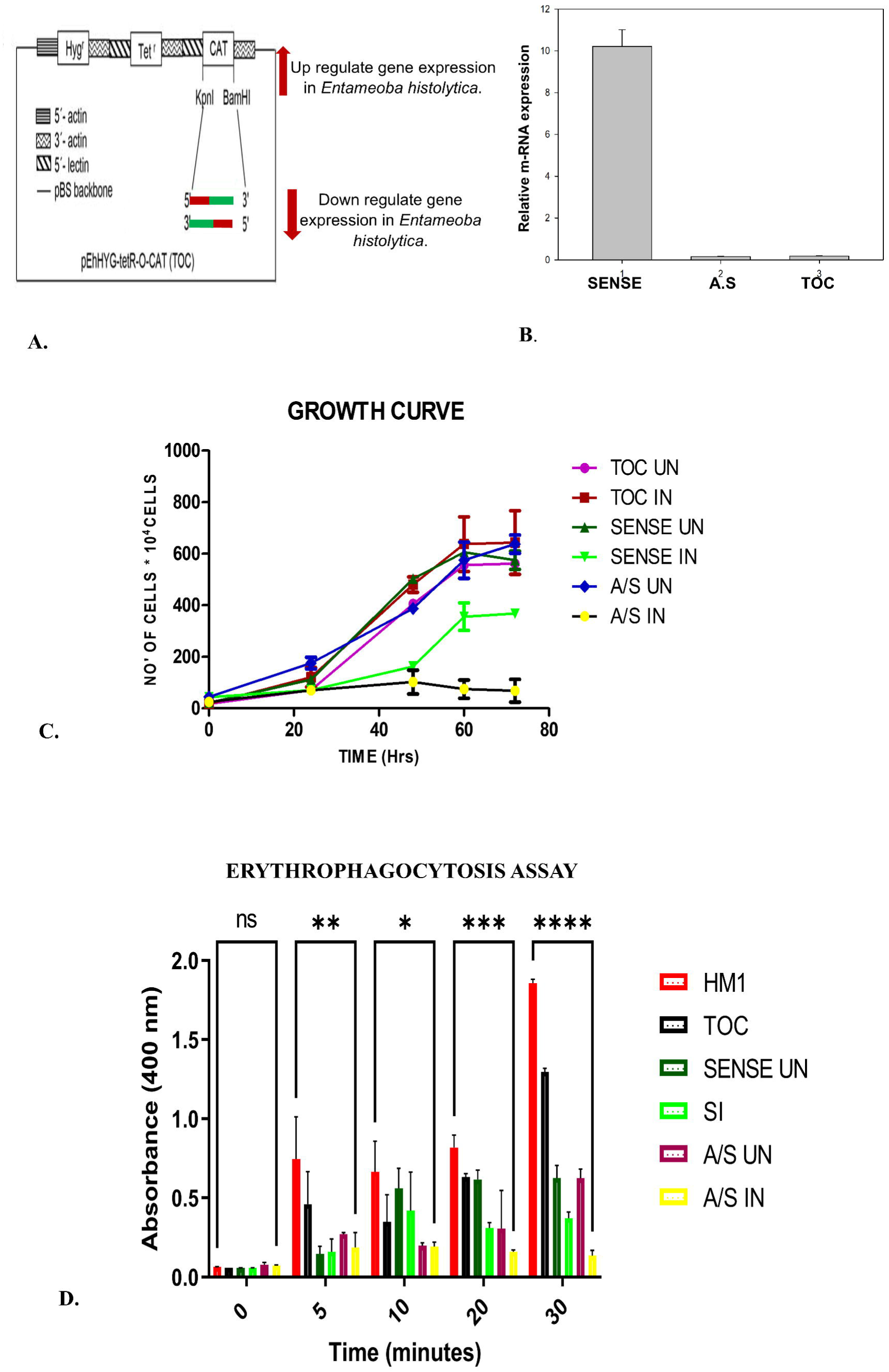
Functional analysis of *Eh*PHDK using tetracycline-inducible expression system in *Entamoeba histolytica*. (**A**.) Vector map of Tet-O-CAT vector (TOC). The *E. histolytica* trophozoite gene *EhPHDK* was cloned in the sense and antisense directions in the TOC vector to examine the effects of upregulation and downregulation. **(B.)** Reverse transcription PCR of the corresponding cDNA, the precise gene suppression was verified. Sense_PHDK (sense), Anti-sense_PHDK (A.S.), Tet-O-CAT vector alone (TOC) shows the relative mRNA expression levels. (C) *Eh*PHDK knockdown severely impairs trophozoite growth: Growth curve of cells of various cell lines when induced with or without 30 ug/ml tetracycline over a period of 72 h. The cell line containing Toc vector with *EhPHDK* cloned in antisense direction shows 90% of growth inhibition in comparison to the control cell lines and cell line containing *EhPHDK* cloned in sense direction. *EhPHDK* plays essential role in the growth of *Entamoeba histolytica* trophozoites. (D) Erythrocyte uptake assay in sense and antisense cell lines. *E. histolytica* cells carrying the above-mentioned constructs were incubated with RBCs for the indicated time points and assessed for RBC uptake by spectrophotometric analysis. The experiments were repeated independently three times in duplicate with error bars indicating the standard error. The statistical comparisons were carried out using one-way AnoVA test (p-value < 0.00005). **(Abbreviation:** S UN-sense uninduced. SI-sense induced, A/S UN-antisense uninduced, A/S IN – anti-sense induced, TOC-Tet-O-CAT vector alone, HM1-wild type HM1: IMSS strain)

For the growth curve study, fresh growth media was inoculated with an equal number of cells. All the cells were synchronised, and released from G0/G1 phase after arrest (S. Fig. 9). At the designated time points, the cells were collected, re-suspended in PBS, and counted on a hemocytometer. The results of three independent tests are presented as mean ± SD (Fig. 8 C). When S_Toc, AS_Toc, and Toc vector alone cell lines were over-expressed with 30 µg/mL of tetracycline, it was shown that the cell line having downregulated PHDK protein expression showed significantly reduced proliferation of amoebic trophozoites by 90%. On the other hand, upregulating the PHDK gene expression in an amoebic cell line also shows declined growth when compared to vector-alone cell strains or uninduced cell strains. Thus, from the growth kinetics experiment, we investigated that optimal gene expression of *EhPHDK* is crucial for cell survivability and proliferation.

By examining the impact of *EhPHDK* downregulation on the rate of phagocytosis, we further illustrated the role of *EhPHDK* in cytoskeleton dynamics and tissue invasion phenomenon. All comparisons were performed using cells that either carried the vector alone, the gene construct without tetracycline, or with a wild-type HM1: IMSS strain of amoeba. RBC uptake decreased by 50% in 30 minutes when *EhPHDK* was overexpressed in the sense cell line. But when the *EhPHDK* gene was overexpressed in anti-sense cell lines, the phagocytosis rate declined up to 90% (Fig. 8 D). That further supports the growth kinetics data, that optimal *EhPHDK* gene activity or protein expression is necessary for controlling cell proliferation, motility, and phagocytosis phenomenon. To further evaluate the experimental data, statistical analysis was done that proves the cell lines chosen for the test have unequal mean i.e., the rate of erythrophagocytosis is significantly different in each of the sample that are selected under one-way Anova test ((p-value <0.00005). Also, the cell lines show unequal variance so this proves that they are independent and each cell lines are drawn from different set of population. All the above findings suggest that *EhPHDK* is a crucial kinase that regulates various cellular responses that determine the survival and pathogenesis of *E. histolytica*.

## DISCUSSION

When a host cell is attached to the amoeba surface, a downstream signalling cascade is activated, causing actin filaments to bind to the plasma membrane and creating the force necessary for the pseudopod to protrude. Phagocytosis is a diverse and multistep process that begins when the particle is attached and terminates soon after phagosomes are formed and separated from the plasma membrane. *E. histolytica* expresses the maximum amount of TMKs in contrast to other protozoan parasites, this parasite has a complex signalling network to perceive its surroundings (1,2,3). The signals that these TMKs gather up are translated by downstream effectors, which may aid the parasite in adapting to the rapidly shifting conditions in the host intestine. Calcium is necessary for the invasive and various endocytic processes, but the molecular mechanism underlying this is still unknown. (16). Since the *E. histolytica* genome codes for 36 CaM-like kinases and 27 Ca^2+^ binding proteins, it is reasonable to assume that these proteins are working together to carry out Ca^2+^-dependent signalling (5). Since amoebae’s cam-like kinases lack a conserved CaM binding site, it is thought that they indirectly regulate Ca^2+^-dependent signalling through CaBPs. (6). According to previous reports, it has been proven that CaBP1 is recruited during erythrophagocytosis and trogocytosis by a C2 domain-containing kinase (protein belongs to the CaM-like kinases family (7). Since CaBP1 is in charge of delivering other proteins to the site of phagocytosis, its recruitment is crucial for the start of the process. The CaBP1-dependent signalling pathway is further extended by another kinase known as *Eh*AK1, an alpha kinase that phosphorylates amoebic actin and recruits the Arp2/3 complex, which is known to be brought in at subsequent steps in a Ca^2+^-dependent manner (10). Arp2/3 complex recruitment is required to initiate actin polymerization at the site of phagocytosis. Actin dynamics appear to depend on the phosphorylation of amoebic G-actin by *Eh*AK1 in vivo as well (11,12).

A number of kinases and phosphatases are activated as a result of adhesion to host cells, aiding in the alteration of phosphoinositide’s found in the amoeba’s plasma membrane (16). Phosphoinositide’s control a number of cellular functions, including membrane trafficking, cytoskeleton reorganization, and endocytosis. PtdIns(3,4,5)P_3_ production is essential for the endocytic process downstream signalling. The trogocytosis-specific AGC family kinase *Eh*AGCK1 is known to be recruited by PtdIns (3,4,5)P_3_ to swallow the living host cells(17). Otherwise, phagocytosis of dead host cells is carried out by a different kinase from the same family, *Eh*AGCK2. (16). 24 AGC family kinases are encoded by the *E. histolytica* genome (8). So, far only two have been identified as being engaged in endocytic activities. It is also known that AGC superfamily kinases affects actin dynamics by acting downstream of PI3K protein (38,39). Although it has been demonstrated that PI3K-PKC activity is necessary for *E. histolytica* to destroy host cells, further proteins in the pathway have not yet been discovered (40). These finding clicks our group interest to identify the role of other proteins that belongs from the AGC superfamily.

Our *Insilico* studies revealed that almost all the multicellular parasites and higher eukaryotes have been found to include PHDK/Akt sequences, frequently in different isoforms. With other parasitic members of the AGC superfamily, the *EhPHDK* had a high degree of sequence identity ranging from 30 to 50%. *EhPHDK* protein shows a maximum 45 % sequence identity with Dictyostelium discoideum serine/threonine kinase. Identity ranged from 33% with (Leishmania infantum, Leishmania panamensis, Leishmania donovani) to 42% with serine/threonine protein kinase from humans. The AlphaFold model suggests that it has three domains i.e., PH domain, Kinase core, and AGC C-terminal domain. Through the predicted structure of the protein, we hypothesize that the PH domain and C-terminal loop may fold in such a way that it masks the kinase core that further controls the phosphorylation activity of the protein.

Numerous findings from our studies, including those based on in-depth live cell imaging, gene silencing, and phagocytosis assay, support the idea that *EhPHDK* is involved in actin-driven activities including phagocytosis and pseudopod extension. From the literature study, it’s already proven that the PH domain of AGC family proteins interacts with very high-affinity PtdIns(3,4,5)P3, which means it’s a membrane-interacting protein (15). The live cell imaging and immunostaining of N-ter GFP tagged *EhPHDK* shows that this protein is ubiquitously dispersed throughout the cell, but during the phagocytosis and other endocytic processes it actively gets recruited to the plasma membrane. *EhPHDK* also participate in fluid-phase micropinocytosis but it’s doesn’t involve in the nibling of live cells. Our immunostaining observations suggest that *EhPHDK* is involved in all the three stages of phagocytosis i.e., initiation, progression / tunnel formation step and also at the tip of the cup during phagosome closure. *EhPHDK* leaves the site as soon the process of phagocytosis completes which means it doesn’t involve in any complete phagosome/ late endosome formation. However, results shown in this report clearly proves that *EhPHDK* actively colocalise with *Eh*CaBP1 during initiation or progression step, hence may play major role in initiating the phagocytosis phenomenon along with *Eh*CaBP1. Our protein-peptide binding study through Bio-layer interferometry reveals the specific amino acid sequence of *EhPHDK* that mimics IQ domain sequences. Moreover, our data suggest that the amino acid sequence (IYKSKITTETVTQKD) lies between the PH domain and a kinase core of the protein, which also serves as a hinge region between these domains and provides the binding site for calcium binding protein1(*Eh*CaBP1). While the other sequences present at the surface of the kinase core don’t show any involvement in the interaction with CaBP’s. From growth curve studies we concluded that when *EhPHDK* gene was supressed in antisense strain, the growth of *E. histolytica’s* suppressed to 80–90%, on the other hand over-expressed cells in sense cell-lines shows 60% of growth inhibition when compared with control cell lines. Thus, these finding clearly reveal that downregulating this gene is fatal for parasites. Also, the optimal *EhPHDK* gene expression is important for controlling various cellular responses like proliferation, motality and growth. Additional proof for this fact is provided by RBC uptake assay, in which similar pattern was observed by overexpressed sense and anti-sense cell lines. When *EhPHDK* gene was upregulated in the sense cell line, RBC uptake dropped by 50% in just 30 minutes. On the other hand, the phagocytosis rate dropped by up to 90% when the *EhPHDK* gene was downregulated in anti-sense cell lines. The crucial role of PH domain containing protein kinases in regulating phagocytosis and motility, determines the survival and pathogenesis of *E. histolytica. All these* finding has major implications in the basic biology of this unicellular parasite as how kinase can remodel cytoskeletal machinery by directly interacting with calcium-binding proteins *Eh*CaBP1 and colocalizing with actin during various endocytic phenomenon. As a result, our findings open the door for further research into the biology of *EhPHDK* in amoebic pathogenesis and show a novel mechanism of modulation of cytoskeletal dynamics in *E. histolytica*.

### Glossary

*EhPHDK* (PH domain containing protein**)-** The designated name for the Akt protein in this study.

## Acknowledgments

We acknowledge the Central Instrument Facility, School of Life Science, JNU, for microscopy studies. We also acknowledge Neha Jawla from National Institute of Immunology, Delhi, India who helped us in FACS experiment. We also thank Dr. Santi Swarup Singha of NII, New Delhi for a generous supply of CHO cells.

## Author Contributions Statement (CRediT)

Shalini Mishra: Led all aspects of the study, including conceptualization, experimentation, analysis, writing (original draft & review and editing). Priya Tomar: Assisted in setting up cell biology experiments. Prof. Alok Bhattacharya: Provided resources and contributed to investigation. Prof. S. Gourinath (Corresponding author): Oversaw all aspects, including conceptualization, methodology, supervision, and manuscript review.

## Funding

The funding is sponsored by STAR award by Science and Engineering Research Board (SERB). SM and PT is thankful to UGC for fellowship. SM is also thankful to ICMR for fellowship.

## Disclosure & Conflict of Interest

The authors declare that the research was conducted in the absence of any commercial or financials relationship that could be constructed as a potential conflict of interest. The authors declare that there is no conflict of interest regarding publication of this paper.

## Ethics Statement

All animal experimentations were performed according to the National Regulatory Guidelines issued by CPSEA (Committee for the Purpose of Supervision of Experiments on Animals), Ministry of Environment and Forest, Govt. of India.

## Notes

### Competing Interest Statement

The authors have declared no competing interest.

