## Supplementary data for "Pleckstrin homology domain-containing serine/threonine kinase plays a crucial role in the survival and phagocytosis of *Entamoeba histolytica*"

Figure S1a

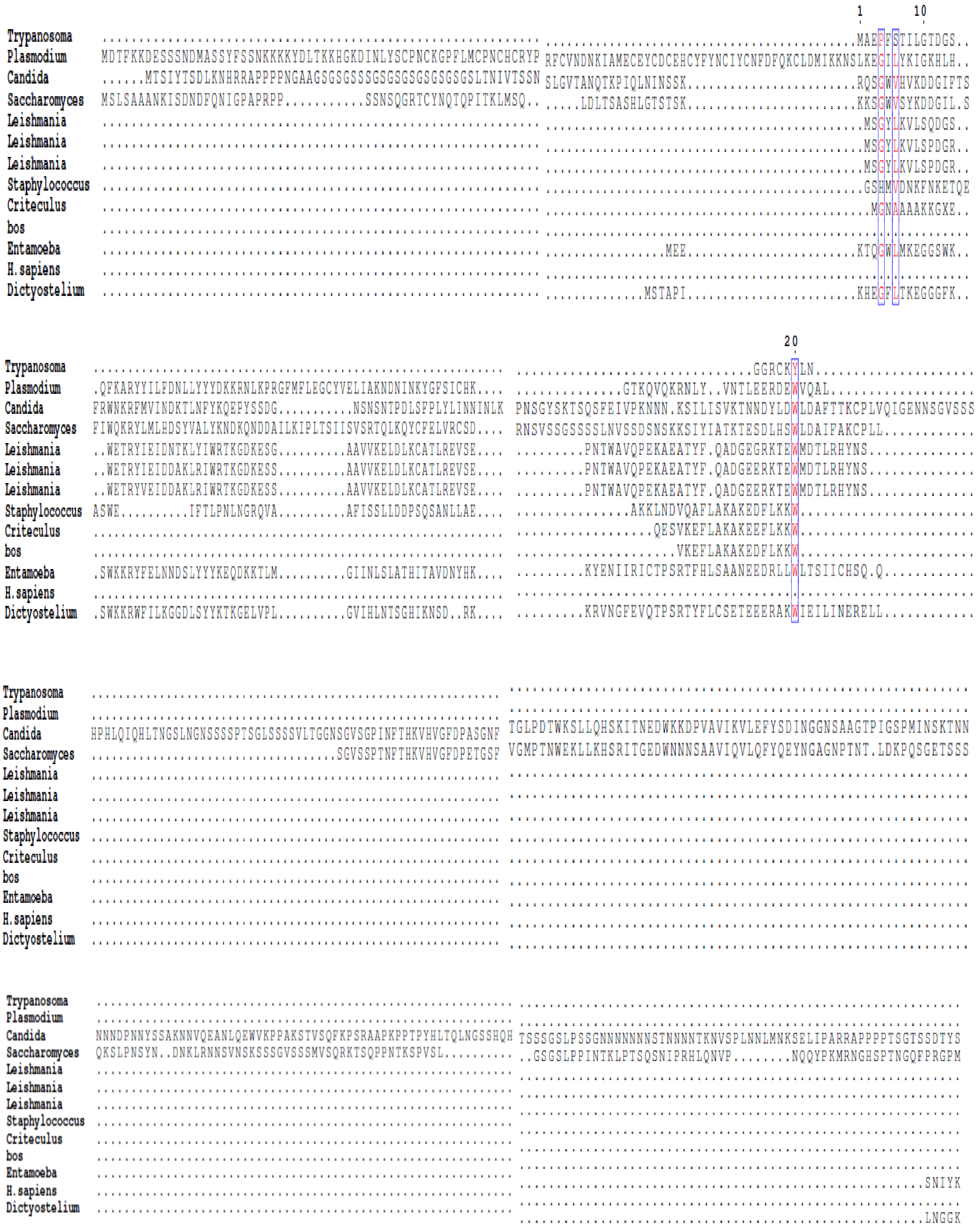
1
10
20
Trypanosoma
Plasmodium
Candida
Saccharomyces
Leishmania
Leishmania
Leishmania
Staphylococcus
Criteculus
bos
Entamoeba
H.sapiens
Dictyostelium

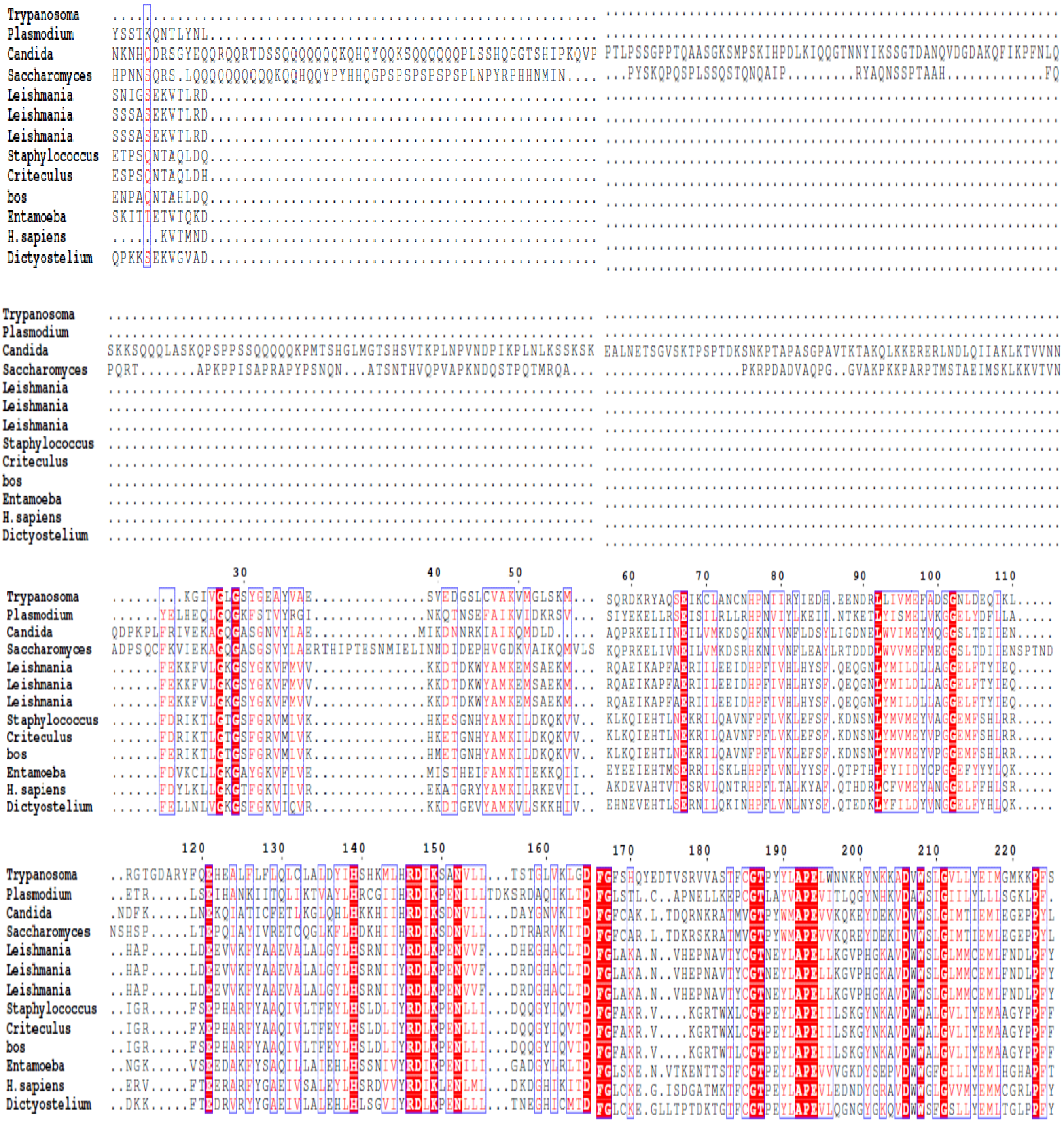
Trypanosoma
Plasmodium
Candida
Saccharomyces
Leishmania
Leishmania
Leishmania
Staphylococcus
Criteculus
bos
Entamoeba
H. sapiens
Dictyostelium
30
40
50
60
70
80
90
100
110
120
130
140
150
160
170
180
190
200
210
220

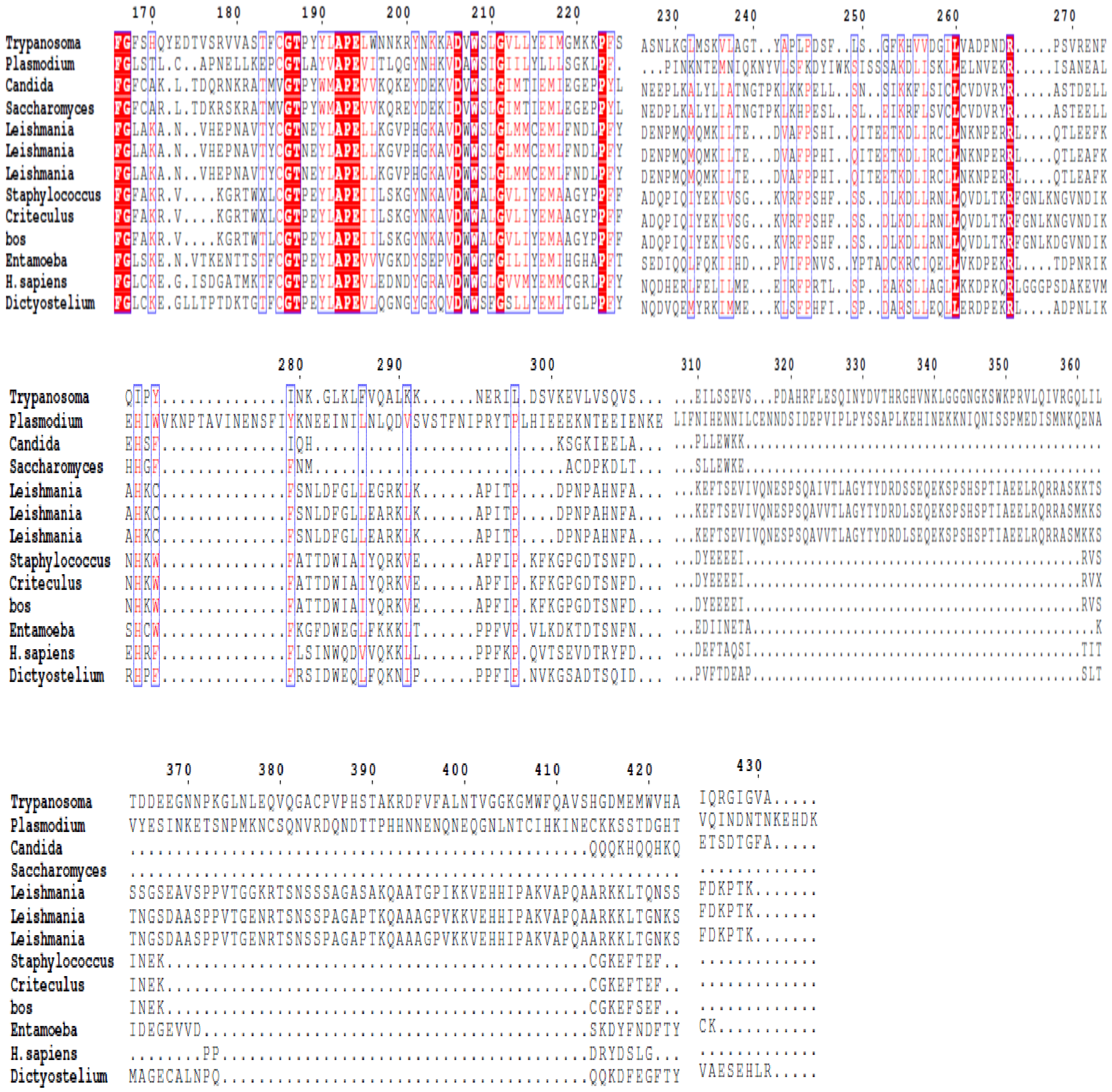

**Supplementary fig1 A. Sequence alignment of EhPHDK proteins**. The full length EhPHDK protein sequences have been aligned with the different parasite, yeast and human EhPHDK isoforms using Clustal W and esprit 3.0 software. That proves only the kinase core is conserved among all the organism, especially among the group of parasites.

Figure S1b

| | | | | | | | | | | | | | |
|---|---|---|---|---|---|---|---|---|---|---|---|---|---|
| 1: AAX79136.1 | 100.00 | 19.33 | 25.16 | 26.30 | 20.19 | 19.71 | 19.71 | 24.61 | 24.68 | 25.99 | 22.73 | 25.75 | 23.17 |
| 2: ETW35934.1 | 19.33 | 100.00 | 21.46 | 20.97 | 19.23 | 20.45 | 20.45 | 24.87 | 28.32 | 28.40 | 26.96 | 27.96 | 28.54 |
| 3: XP_723573.1 | 25.16 | 21.46 | 100.00 | 45.88 | 22.02 | 21.76 | 21.76 | 23.03 | 23.17 | 23.92 | 22.78 | 26.33 | 23.08 |
| 4: KAF1902903.1 | 26.30 | 20.97 | 45.88 | 100.00 | 22.31 | 22.31 | 22.31 | 22.86 | 23.95 | 25.17 | 21.76 | 27.64 | 24.23 |
| 5: AKE32416.1 | 20.19 | 19.23 | 22.02 | 22.31 | 100.00 | 93.73 | 93.53 | 32.89 | 34.50 | 36.39 | 33.89 | 38.89 | 34.76 |
| 6: XP_003862798.1 | 19.71 | 20.45 | 21.76 | 22.31 | 93.73 | 100.00 | 99.80 | 32.89 | 35.09 | 36.39 | 33.17 | 39.22 | 35.00 |
| 7: XP_001466930.1 | 19.71 | 20.45 | 21.76 | 22.31 | 93.53 | 99.80 | 100.00 | 32.89 | 35.09 | 36.39 | 33.17 | 39.22 | 35.00 |
| 8: pdb\|5X3F\|B | 24.61 | 24.87 | 23.03 | 22.86 | 32.89 | 32.89 | 32.89 | 100.00 | 93.16 | 96.43 | 35.16 | 40.45 | 37.50 |
| 9: pdb\|5NW8\|A | 24.68 | 28.32 | 23.17 | 23.95 | 34.50 | 35.09 | 35.09 | 93.16 | 100.00 | 96.43 | 39.13 | 40.78 | 42.07 |
| 10: pdb\|2UZT\|A | 25.99 | 28.40 | 23.92 | 25.17 | 36.39 | 36.39 | 36.39 | 96.43 | 96.43 | 100.00 | 40.30 | 40.45 | 43.07 |
| 11: XP_656547.1 | 22.73 | 26.96 | 22.78 | 21.76 | 33.89 | 33.17 | 33.17 | 35.16 | 39.13 | 40.30 | 100.00 | 42.02 | 45.45 |
| 12: pdb\|1GZK\|A | 25.75 | 27.96 | 26.33 | 27.64 | 38.89 | 39.22 | 39.22 | 40.45 | 40.78 | 40.45 | 42.02 | 100.00 | 52.41 |
| 13: XP_646888.1 | 23.17 | 28.54 | 23.08 | 24.23 | 34.76 | 35.00 | 35.00 | 37.50 | 42.07 | 43.07 | 45.45 | 52.41 | 100.00 |

**Supplementary fig1 B.** Percentage identity matrix with other orthologs was calculated by Clustal2.1 (the serial number follow 1-13 are the accession numbers of AGC proteins from different organism sequentially listed in the alignment result)

**1.** Trypanosoma brucei AGC **2.** Plasmodium AGC **3.** Candida albicans AGC **4.** S. Cerevisiae AGC **5.** Leishmania panamensis AGC **6.** L.donovani AGC **7.** L.infantum AGC **8.** Staphylococcus aureus AGC **9.** Cricetulus griseus AGC **10.** Bos tarus AGC **11.** Entamoeba histolytica AGC 12. Human AGC 13. Dictyostelium discoideum AGC.

Figure S2

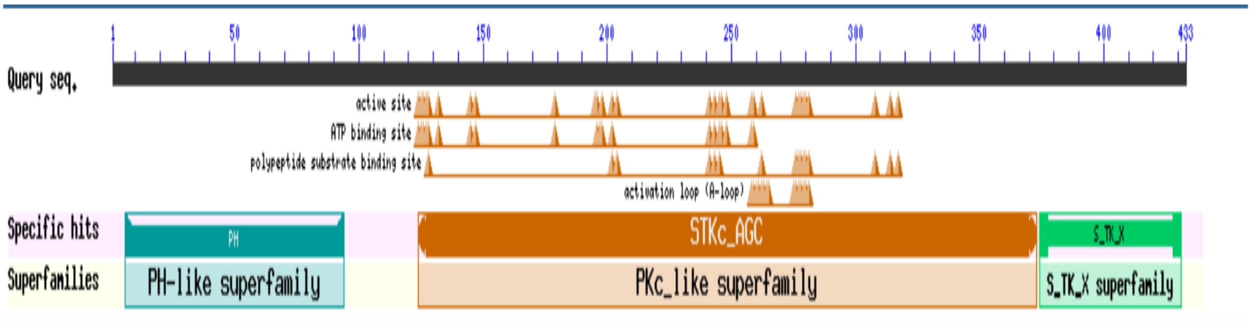


**Supplementary fig2.**

Protein model of *Eh*PHDK ( Protein_id ="XP_656547.1"/ EHI_042150 constructed through conserved domain database of NCBI showing active site, ATP binding site and polypeptide binding site.

Figure S3

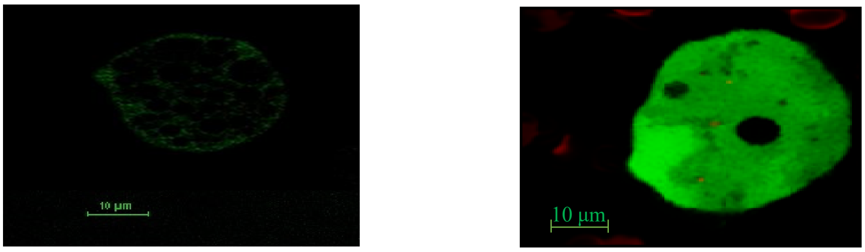


A.

**HM1: IMSS trophozoite cell** **GFP-EhPHDK expressing cells**

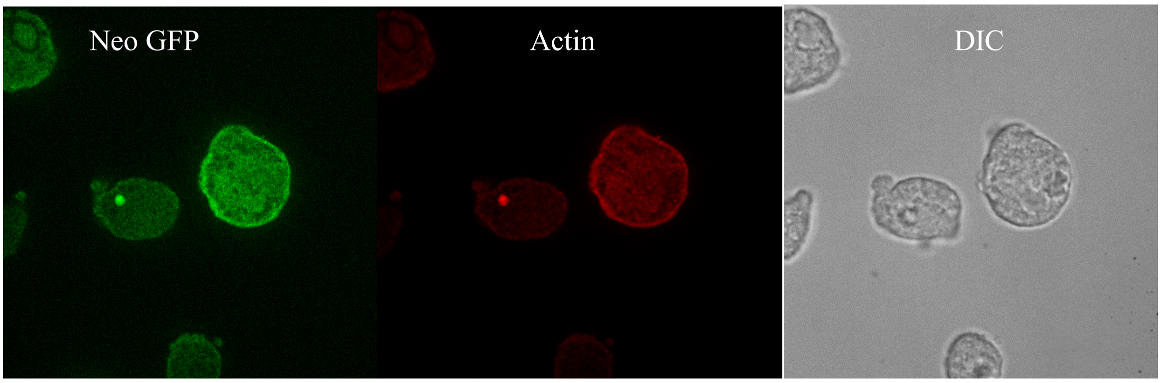


**B. Immunostaining image of neo-GFP transfected trophozoite cells**

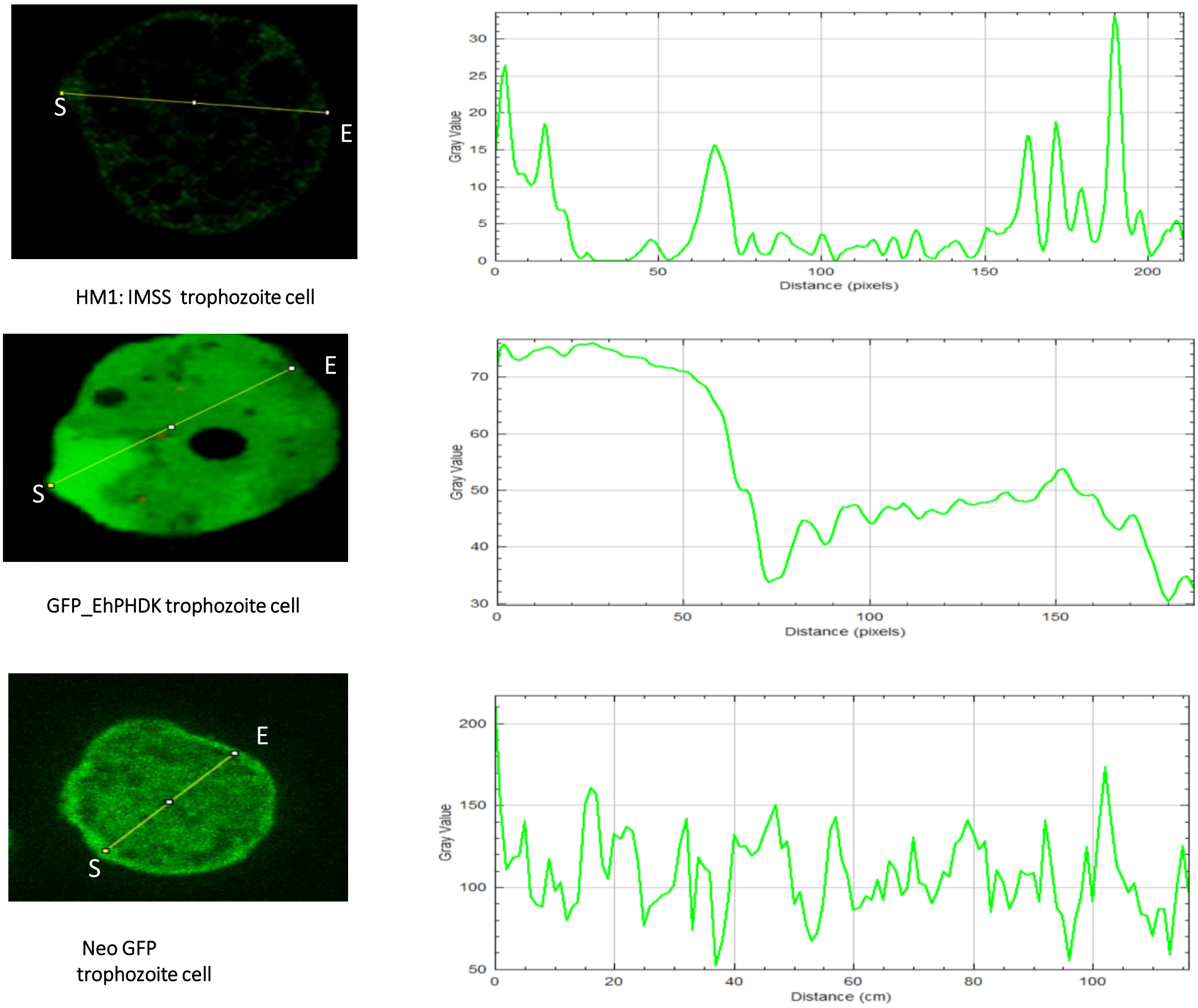


C.

**Supplementary fig3. A.** Live-cell image comparing fluorescence intensity between WT trophozoite (unstained, non-transfected) and GFP-tagged EhPHDK-transfected trophozoite. **B.** Immunostained Neo-GFP transfected trophozoite image showing **green (neo-GFP), red (Actin), and differential interference contrast (DIC). C.** Fluorescence intensity profile across the cell, measured from the pseudopod to the opposite side. The graph compares fluorescence signals in HM1 (wild-type control), neo GFP (control for general GFP expression), and GFP-PHDK. The neo GFP control exhibits uniform fluorescence intensity across the cell, while HM1 shows negligible fluorescence, confirming no GFP expression in wild-type cells. In contrast, GFP-PHDK fluorescence is significantly higher near the pseudopod region, indicating preferential localization of PHDK protein. These results suggest a potential role of PHDK in pseudopod formation, cellular motility, or cytoskeletal dynamics.

Figure S4

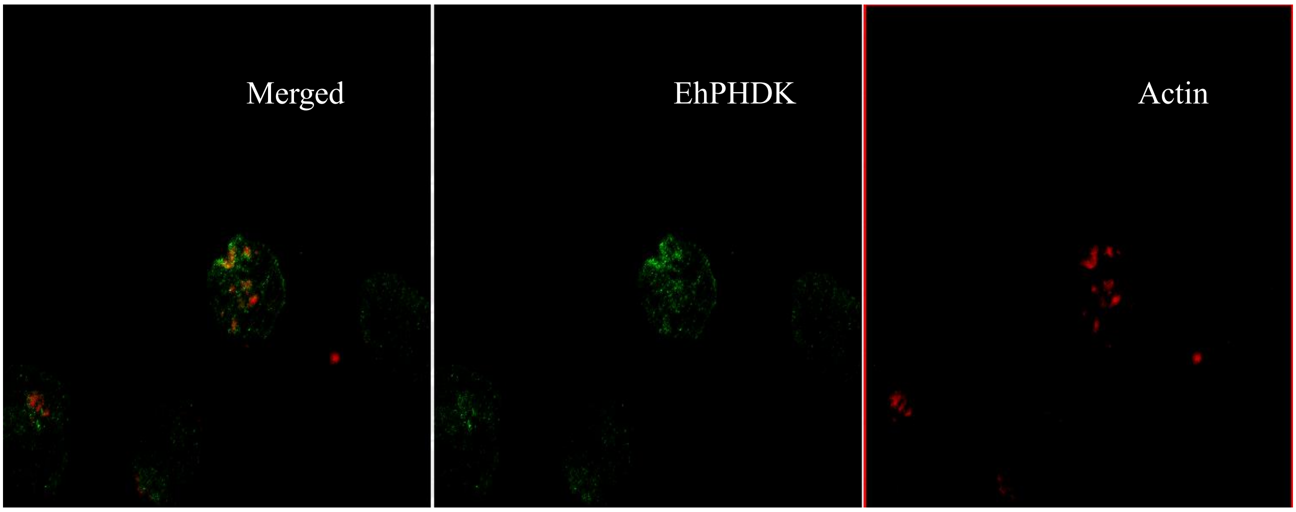


A

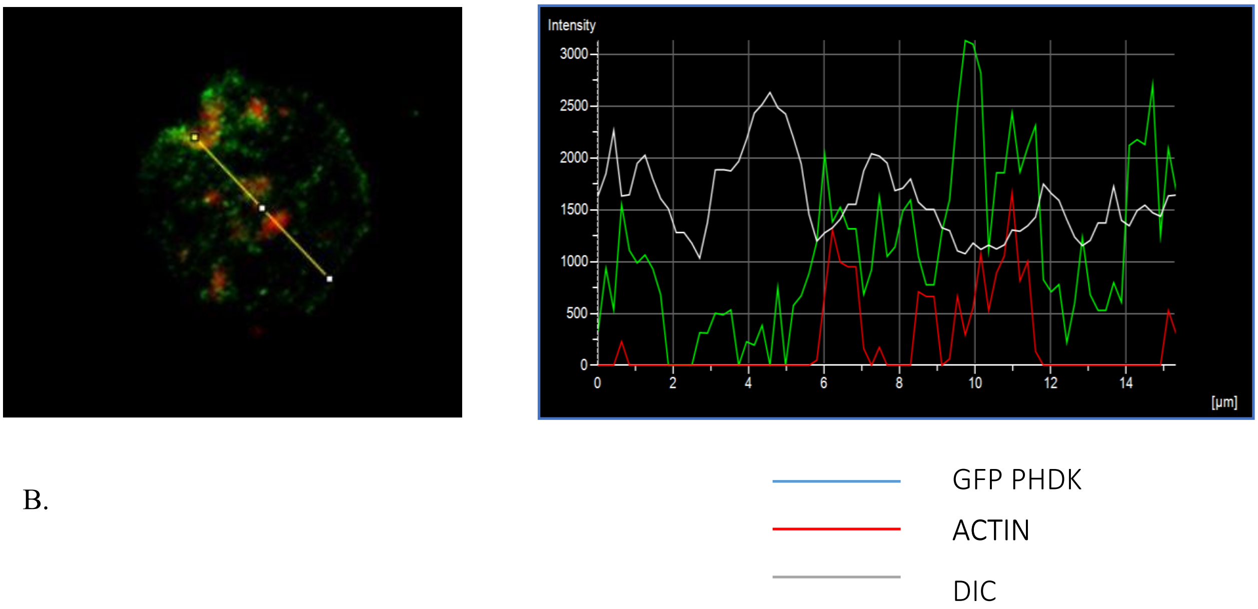


B.

**Supplementary fig4. Co-localization of GFP_EhPHDK along with actin in the phagocytic cup:** Immunostaining of N-ter GFP fused *Eh*PHDK suggest that it actively involved during pseudopod formation and regulate the actin polymerisation in *E. histolytica* **(correlation coefficient : 0.784 )**

Figure S5

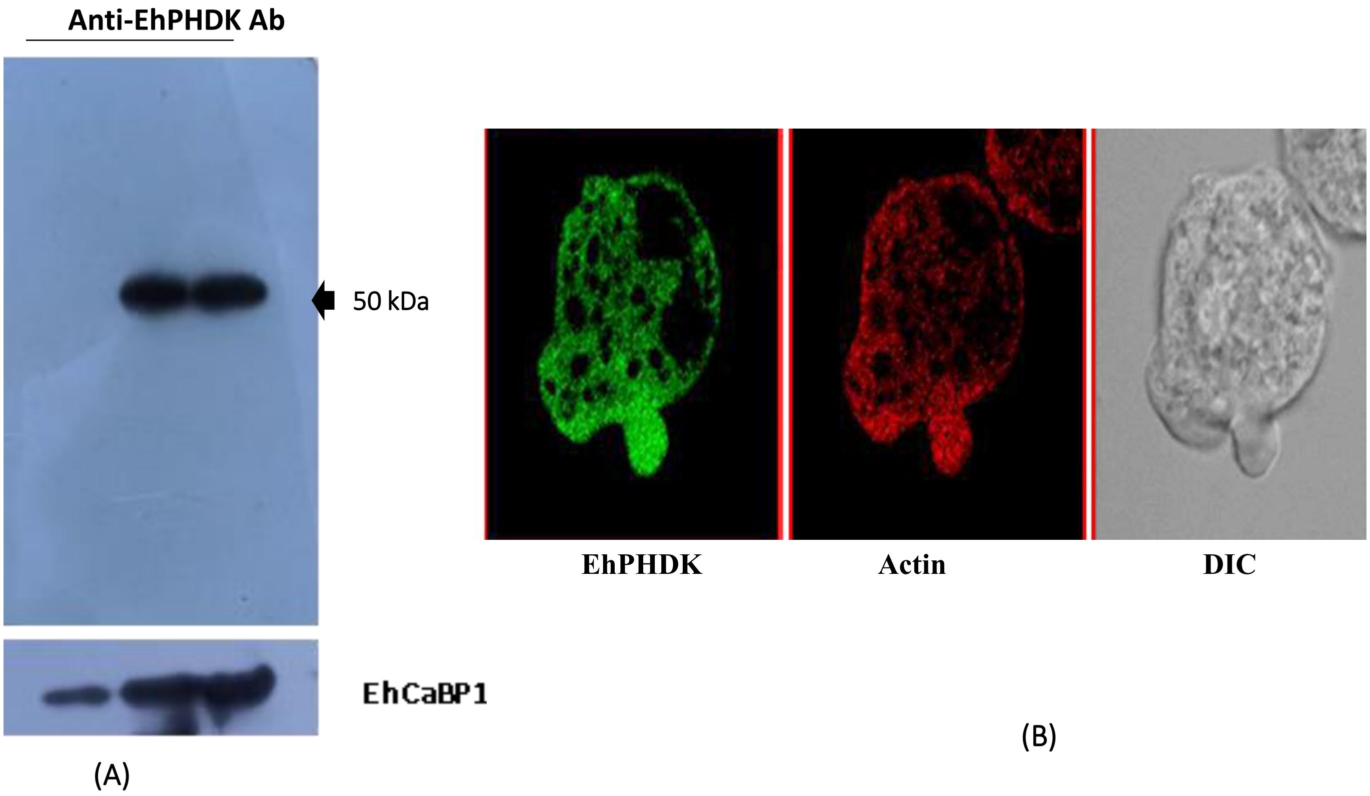


**Supplementary fig 5. Western blot and IHC using Anti-EhPHDK antibody production. (A)** Western blot analysis was used to determine the specificity of the anti-EhPHDK antibody produced against recombinant protein. Entamoeba lysate (50 150µg,150µg,) was probed with anti-EhPHDK (1:1,500). EhPHDK's molecular weight was predicted to be 50 kDa. EhCaBP1 was used as the loading control. **(B).** Localization and expression pattern were also observed when we carried out immunostaining of wild-type (HM1: IMSS) cells with an *Eh*PHDK antibody. The results predicts that GFP fused protein shows the similar localization pattern as the native protein shows.

Figure S6

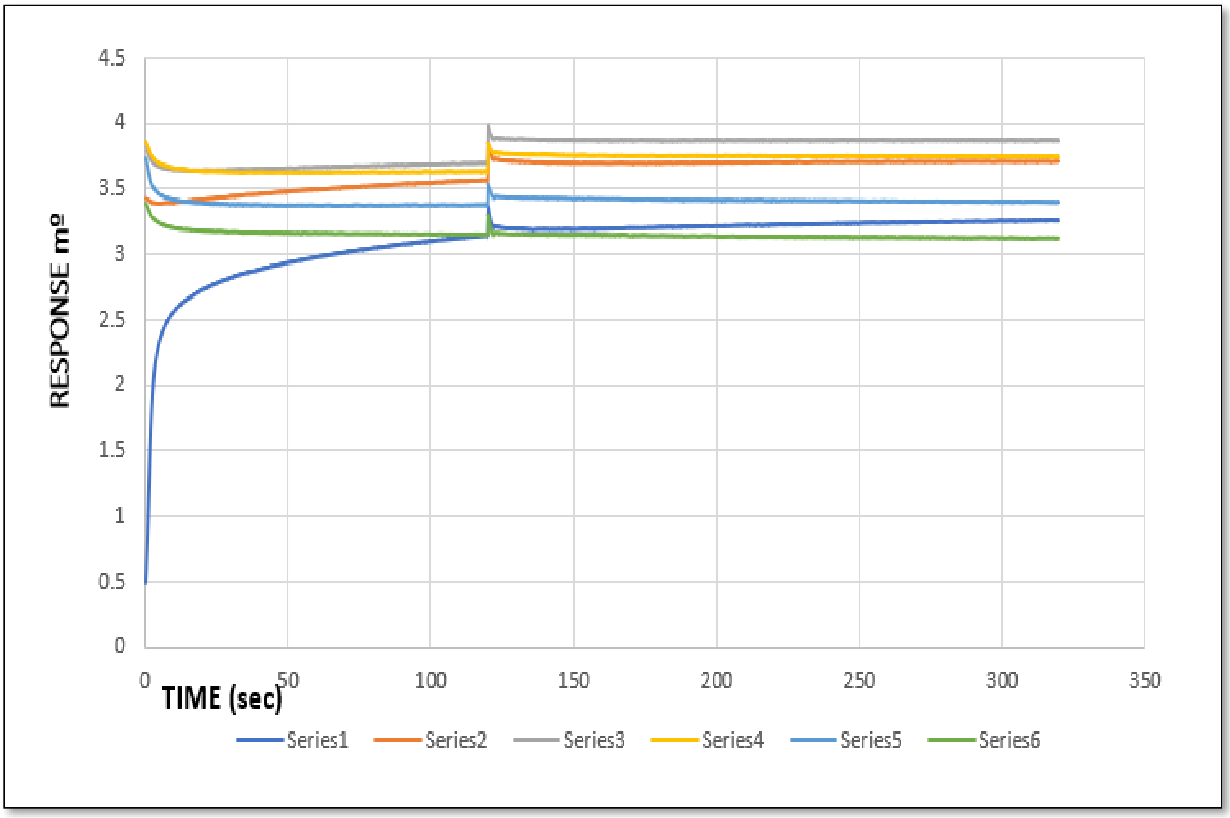


**Supplementary fig 6.** Bio-layer interferometry graph showing the binding affinity of EhPHDK Peptide1 for EhCaBP1(overlayed on Ni-NTA chip. The colours represent different peptide 1 concentrations, ranging from (300 µM- 550 µM /series1- series6) exhibits no affinity for EhCaBP1. The Kd value is 0.000001 µM µM calculated from graph.

Figure S7

Merge EhCoactosin EhPHDK Actin

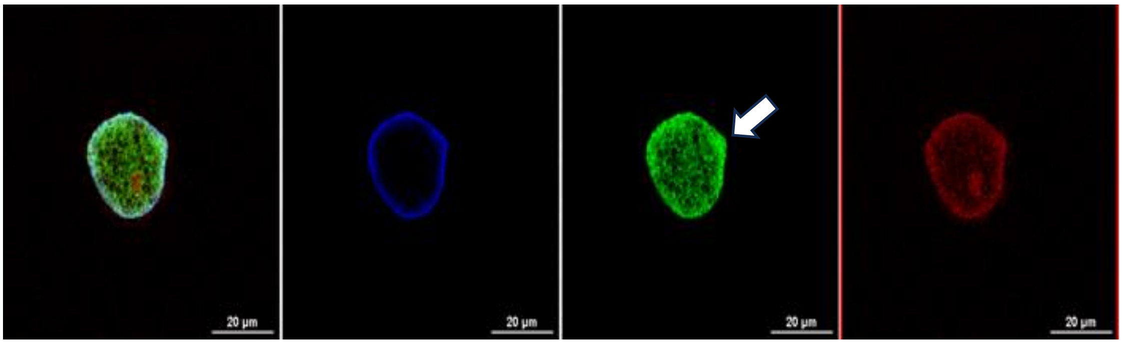


A

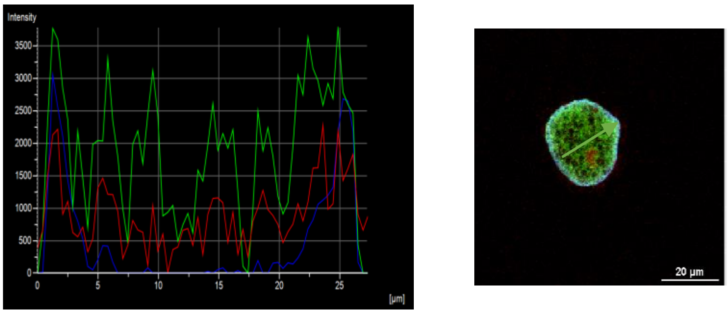


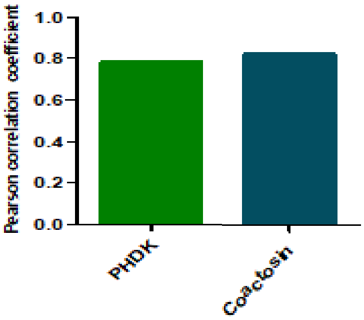


B

C

**Supplementary fig 7. (A.)** Immunostaining of N-ter GFP tagged *Eh*PHDK suggest that it actively involved during the phagocytic cup initiation and actively colocalize with Coactosin. **(B.)** The intensity of GFP_PHDK along the arrow shown in micrograph at different points. The intensity profile at budding pseudopod formed by the trophozoite show enrichment along the line of analysis and also show its localization with EhCoactosin (Pearson's correlation 0.7075). **(C)** Pearson correlation coefficient value were calculated using NIS element software by selecting area enclosing the phagocytosis cup, at the base and also the tip of endocytic cups from different trophozoites. Statistical analysis was performed by using graph pad prism software.

Figure S8

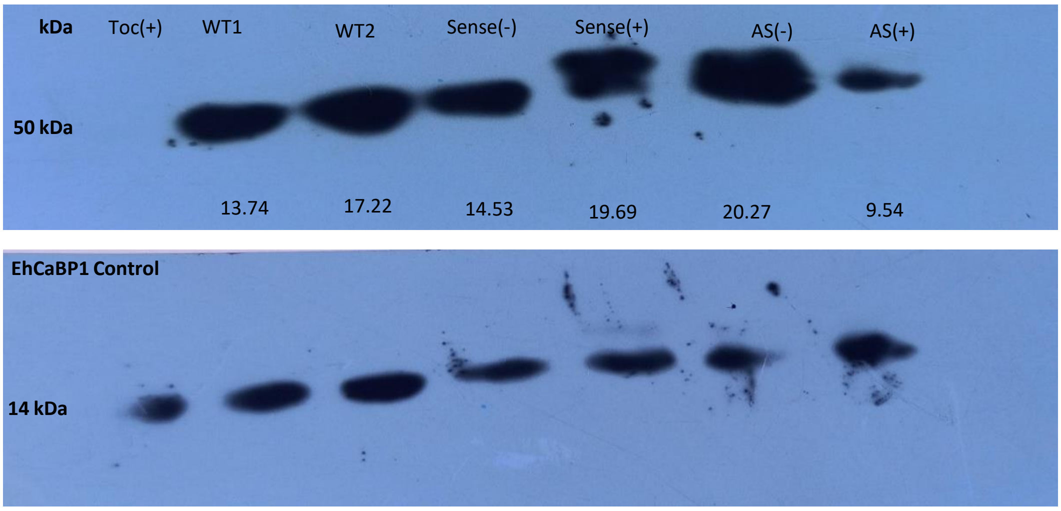


**Toc**- Control cell with only vector
**Sen**- Upregulated cell lines created with transfection of Entamoeba cells with Tet-O-CAT plasmid having gene PHDK in sense direction
**AS**- Downregulated cell lines created with transfection of Entamoeba cells with Tet-O-CAT plasmid having gene PHDK in antisense direction
**WT-** HM1:IMSS cell lysate.

- - Indicates not induced cell lines
- + Indicates induced cell lines with 30µg/ml hygromycin

**Supplementary fig 8.** Western blotting. Protein levels were determined by western blotting in the indicated cells. '+' indicates tetracycline addition (30 µg·ml-1 ) for 48 h and '- ' indicates no tetracycline induction. 150 µg of whole-cell lysates were separated on SDS-PAGE and transferred to PVDF membrane, which was then probed with the rabbit anti-EhPHDK antibody (1:1500) followed by detection with HRPO conjugated anti-rabbit secondary antibody (1:10000). EhCaBP1 was used as the loading control. Band intensity in each lane was determined by performing densitometry using ImageJ software.

Figure S9

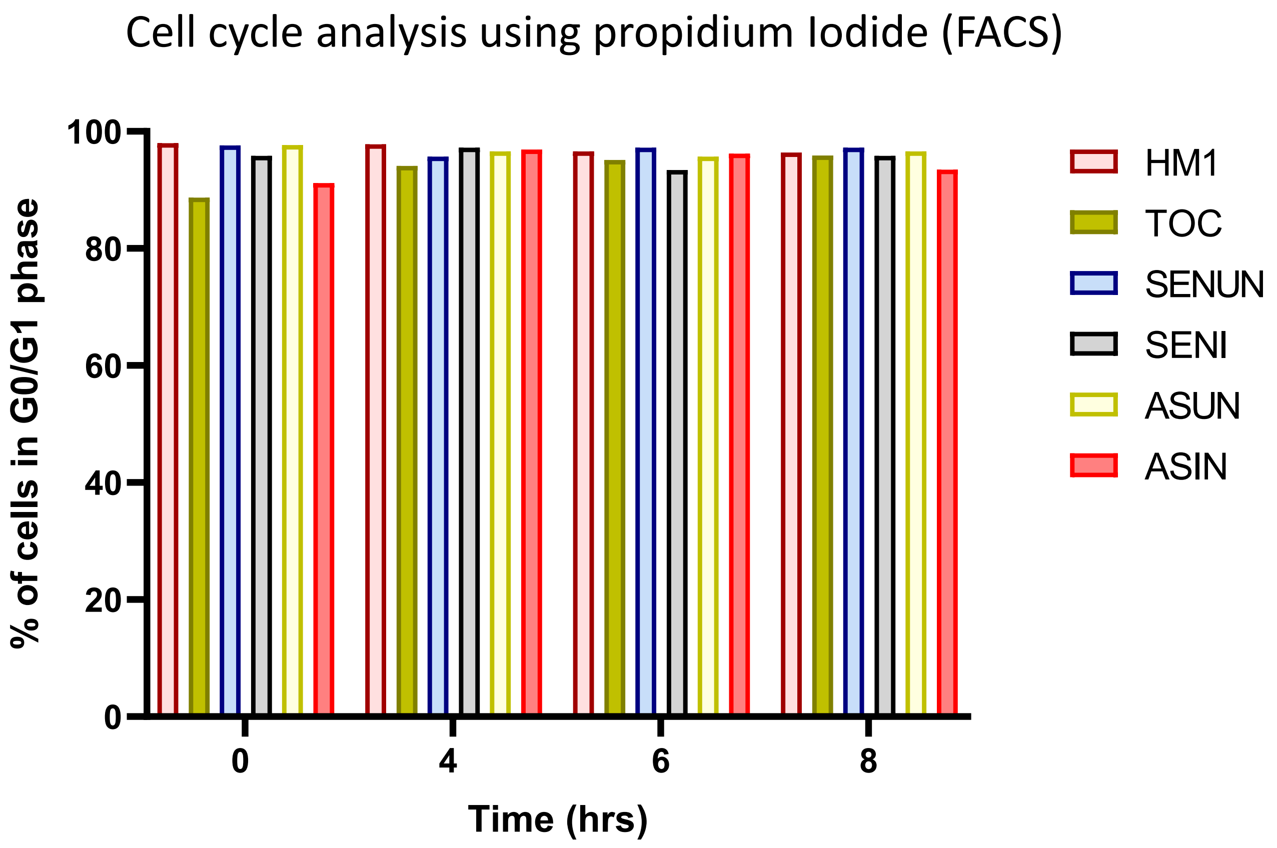


Cell cycle analysis using propidium Iodide (FACS)
Shows more then 90% of cells are in same phase (G0/G1 stage)

Supplementary fig 9

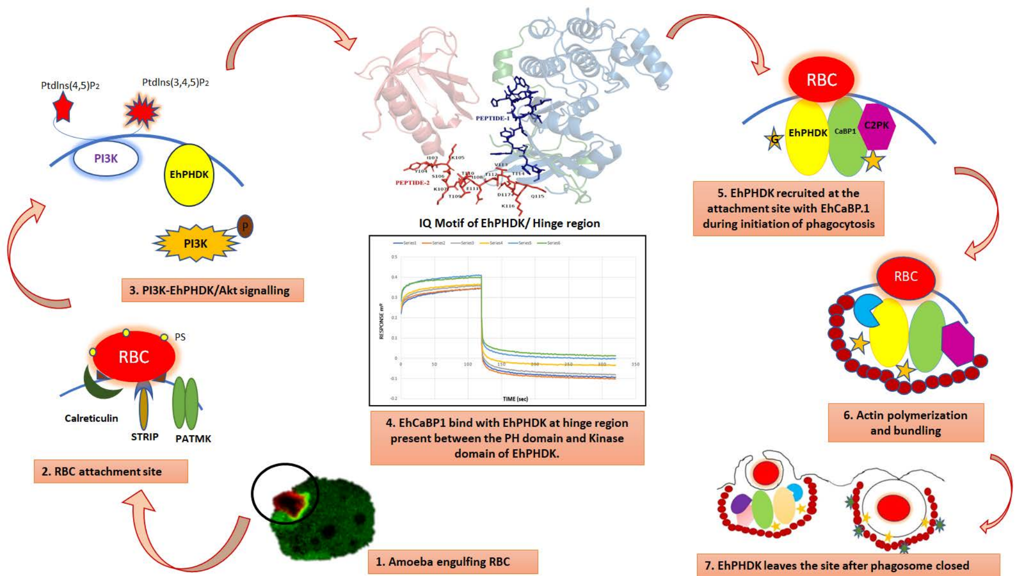

PtdIns(4,5)P2
PtdIns(3,4,5)P2
PI3K
EhPHDK
P
PI3K
3. PI3K-EhPHDK/Akt signalling
PEPTIDE-1
PEPTIDE-2
IQ Motif of EhPHDK/ Hinge region
RESPONSE m°
TIME (sec)
4. EhCaBP1 bind with EhPHDK at hinge region present between the PH domain and Kinase domain of EhPHDK.
RBC
G
EhPHDK
CaBP1
C2PK
5. EhPHDK recruited at the attachment site with EhCaBP.1 during initiation of phagocytosis
RBC
6. Actin polymerization and bundling
PS
RBC
Calreticulin
STRIP
PATMK
2. RBC attachment site
1. Amoeba engulfing RBC
7. EhPHDK leaves the site after phagosome closed
